# Impaired maturation of wild-type superoxide dismutase 1 associated with neurodegeneration in Parkinson disease brain and a novel murine model

**DOI:** 10.1101/2024.10.29.620972

**Authors:** Amr H. Abdeen, Benjamin G. Trist, Sara Nikseresht, Richard Harwood, Stéphane Roudeau, Benjamin D. Rowlands, Fabian Kreilaus, Veronica Cottam, David Mor, Miriam Richardson, Joel Siciliano, Julia Forkgen, Greta Schaffer, Sian Genoud, Anne A. Li, Nicholas Proschogo, Bernadeth Antonio, Gerald Falkenberg, Dennis Brueckner, Kai Kysenius, Jeffrey R. Liddell, Sandrine Chan Moi Fat, Sharlynn Wu, Jennifer Fifita, Thomas E. Lockwood, David P. Bishop, Ian Blair, Richard Ortega, Peter J. Crouch, Kay L. Double

## Abstract

Atypical wild-type superoxide dismutase 1 (SOD1) protein misfolding and deposition occurs specifically within the degenerating substantia nigra pars compacta (SNc) in Parkinson disease. Mechanisms driving the formation of this pathology and relationship with SNc dopamine neuron health, are yet to be fully understood. We applied proteomic mass spectrometry and synchrotron-based biometal quantification to post-mortem brain tissues from the SNc of Parkinson disease patients and age-matched controls to uncover key factors underlying the formation of wild-type SOD1 pathology in this disorder. We then engineered brain copper deficiency and upregulated SOD1 protein levels in a novel mouse strain, termed the SOCK mouse, to verify their involvement in the development of Parkinson-like wild-type SOD1 pathology and their impact on dopamine neuron health. Soluble SOD1 protein in the degenerating Parkinson disease SNc exhibited altered post-translational modifications, which may underlie changes to the enzymatic activity and aggregation of the protein in this region. These include decreased copper binding, dysregulation of physiological glycosylation, and atypical oxidation and glycation of key SOD1 amino acid residues. We demonstrated that the biochemical profile introduced in SOCK mice promotes the same post-translational modifications and the development of Parkinson-like wild-type SOD1 pathology in the midbrain and cortex. This pathology accumulates progressively with age and is accompanied by nigrostriatal degeneration and dysfunction, which occur in the absence of α-synuclein deposition. These mice do not exhibit weight loss nor spinal cord motor neuron degeneration, distinguishing them from transgenic mutant SOD1 mouse models. This study provides the first *in vivo* evidence that mismetallation and altered post-translational modifications precipitates wild-type SOD1 misfolding, dysfunction and deposition in the Parkinson disease brain, which may contribute to SNc dopamine neuron degeneration. Our data position this pathology as a novel drug target for this disorder, with a particular focus on therapies capable of correcting alterations to SOD1 post-translational modifications.

## Introduction

Parkinson disease is the fastest growing neurological disorder worldwide [1, 2], with a global prevalence of over 8 million individuals that is expected to double by 2050 [3]. The availability of disease-modifying treatments will be critical to effectively manage this disorder, however to date, no such interventions have been approved for clinical use. Movement dysfunction in Parkinson disease primarily results from the progressive death of dopamine-producing neurons in the substantia nigra pars compacta (SNc) [4, 5], which is thought to result from a complex confluence of molecular changes. Amongst these, the abnormal misfolding and deposition of key cellular proteins represents a driving factor, with atypical α-synuclein protein constituting a focal point in the search for protein-based biomarkers [6] and disease-modifying therapies [7]. However, despite a clear role for α-synuclein in Parkinson disease, deposition of α-synuclein is not present in all Parkinson disease patients [8], nor is it restricted to degenerating brain regions [9, 10]. It is therefore likely that additional deleterious molecular pathways contribute to the selective vulnerability of SNc dopamine neurons in this disorder, suggesting exploration of complementary disease targets and approaches is required.

Our group discovered the accumulation of structurally-disordered, immature forms of another protein, superoxide dismutase 1 (SOD1), in post-mortem brain tissues from idiopathic Parkinson disease patients [11]. These alterations are not associated with *SOD1* gene mutations [12], unlike well-documented mutant SOD1 pathology underlying neurodegeneration in rare inherited forms of another neurological disorder, amyotrophic lateral sclerosis (ALS) [13]. Wild-type disordered (dis)SOD1 is reported to compromise neuron health *in vitro* [14] and is restricted to degenerating regions of the Parkinson disease brain, including the SNc, where it is found during the earliest (pre-clinical) stages of this disorder when cell death is most rapid [11]. Collectively, these findings indicate that wild-type disSOD1 protein may be associated with the selective death of SNc dopamine neurons in Parkinson disease. Here we characterize atypical molecular changes in SOD1 in the human Parkinson disease brain and demonstrate that the development of wild-type disSOD1 in the SNc results in progressive age-dependent dopamine neuron death in a novel mouse model expressing this pathology. Our data suggest the development of disSOD1 in the SNc plays a role in Parkinson disease etiology and may represent a novel drug target for modifying the course of Parkinson disease.

## Results

### Post-translational modification of SOD1 is altered specifically in the degenerating Parkinson disease SNc

Superoxide dismutase 1 is a ubiquitously-expressed antioxidant protein that is responsible for detoxifying damaging superoxide radicals produced by mitochondrial respiration and other essential cellular processes [14]. Nascent, unfolded human SOD1 is structurally disordered and lacks antioxidant activity, while mature SOD1 constitutes one of the most thermodynamically-stable cytosolic proteins [14, 15]. Maturation of the immature protein to enzymatically-active, mature SOD1 occurs via a series of stabilizing post-translational modifications (PTMs), including the binding of two copper and two zinc ions [14]. We have previously demonstrated an accumulation of immature disSOD1 conformers in the post-mortem SNc of idiopathic Parkinson disease patients [11], which we posit reflects decreased SOD1 copper binding consistent with our observation of a 65% decrease in copper within this brain region [16]. Here, we quantified metals bound to SOD1 dimers isolated from post-mortem brain tissues of Parkinson disease patients and age-matched controls (**Fig. 1a**; **Supplementary Tables 1, 2**), in whom the absence of *SOD1* gene mutations had been confirmed (**Supplementary Table 3**). Metallation analyses were performed according to our published method [17, 18], which combines size-exclusion chromatography and native isoelectric focusing to enrich soluble SOD1 protein >99-fold in human post-mortem tissue extracts. SOD1-bound copper and zinc are then quantified using highly-sensitive synchrotron X-ray fluorescence (SXRF) analysis. Dimeric SOD1 purified from both the SNc and occipital cortex (OCx) of control cases contained copper and zinc in an almost exact 1:1 ratio, consistent with the theoretical ratio of these elements bounds to mature SOD1 [17]. By contrast, this ratio was significantly lower (0.83 copper: 1 zinc) in the degenerating SNc of Parkinson disease cases compared with controls, but was unchanged in the OCx, a non-degenerating brain region in Parkinson disease (1.08 copper: 1 zinc)(**Fig. 1a**). This indicates decreased copper binding to SOD1 specifically within the Parkinson disease SNc, which we speculate results from the combination of lower bioavailable copper [16] and higher SOD1 protein expression [11] we reported within this region. These data align with the copper:zinc ratio reported within SOD1 aggregates in the Parkinson disease SNc [19], supporting a mechanism of SOD1 self-assembly driven by metal-deficient soluble protein isoforms.

**Fig. 1.**
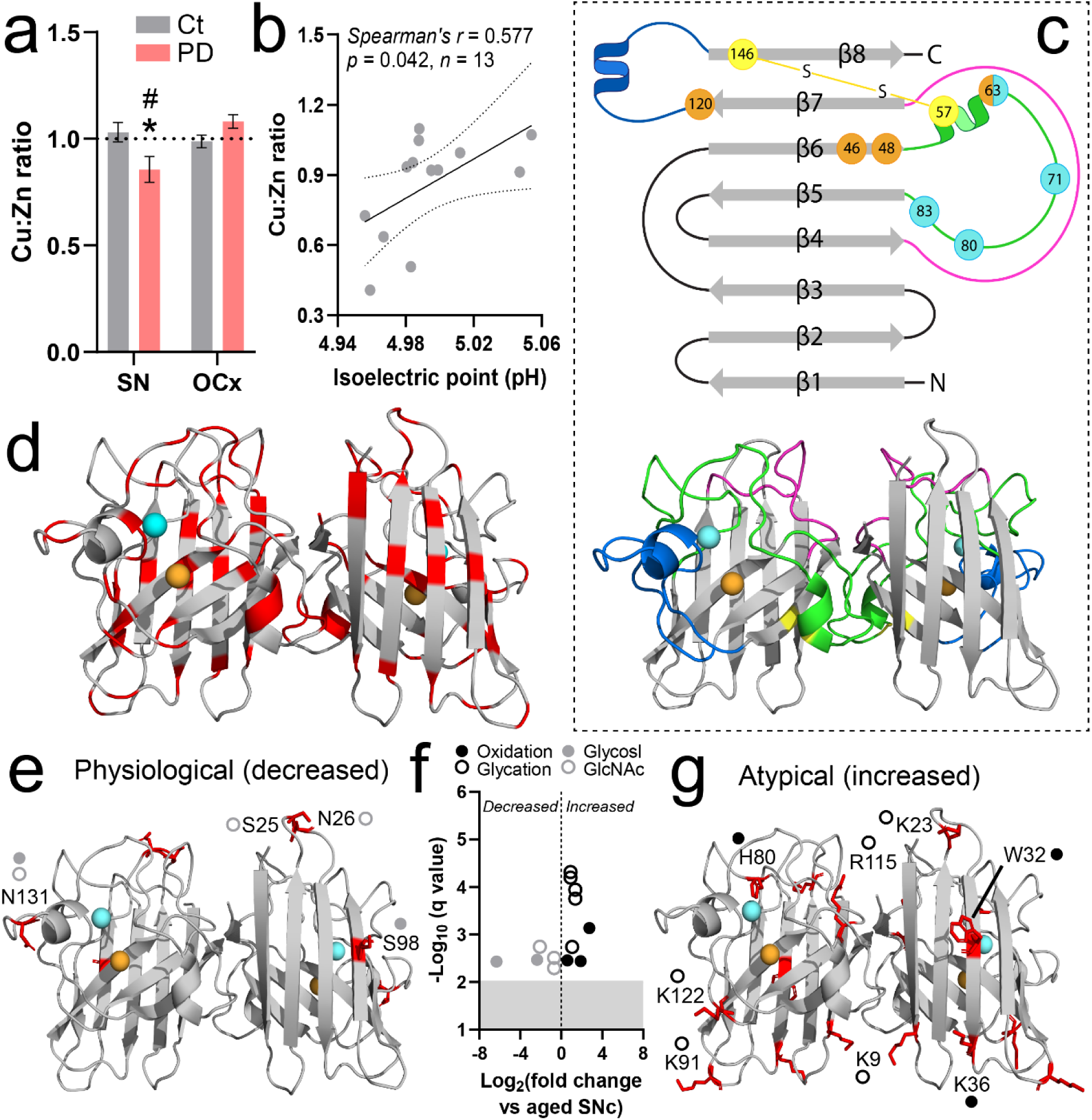
Altered SOD1 post-translational modification in the post-mortem Parkinson disease SNc. **a.** The ratio of Cu:Zn bound to SOD1 was decreased in the post-mortem SNc of Parkinson disease (PD) patients (*n* = 14) compared with controls (*n* = 14), but was unchanged in the PD OCx (*n* = 17) compared with controls (*n* = 18)(Two-way ANOVA: *p* = 0.0039, F = 9.22; Sidak’s multiple comparisons post-hoc tests: \**p =* 0.022 for PD vs Ct SNc, *p* = 0.24 for PD vs Ct OCx, #*p* = 0.0009 for PD SNc vs PD OCx). Data represents mean ± SEM. **b.** Lower SOD1 Cu:Zn ratios were correlated with decreased SOD1 isoelectric point in the PD SNc. Spearman’s *r* coefficient, the *p* value and the number of XY pairs analyzed (*n*) are stated within the panel. **c.** Mature SOD1 is dimeric, with each monomer comprising an eight-stranded β-barrel (grey) that binds one Cu (orange) and Zn (cyan) ion. The electrostatic loop (blue) guides anionic superoxide towards Cu in the active site using a series of charged and polar residues. Zinc coordination is facilitated by three histidine residues and one aspartic acid residue (cyan) within the metal-binding loop (green), whilst copper coordination is mediated by four histidine residues (orange). The disulfide loop (yellow) is a substructure within the metal-binding loop, containing one of two cysteine residues that form an intramolecular disulfide bond within SOD1 protein (yellow). The Greek key loop (pink) forms a plug at one pole of the β-barrel and contributes to dimer interface stability. **d.** Distribution of all residues identified as sites of PTMs in SOD1 protein isolated from the SNc of PD cases and controls (highlighted in red, residues listed in **Supplementary Table 4**). **e.** Distribution of residues exhibiting significantly lower levels of physiological modifications (glycosylation, acetylglucosamination). **f.** A significant decrease in glycosylation (glycosyl) of S98 and N131 and acetylglucosamination (GlcNAc) of S25, N26 and N131 was identified in the SNc of PD patients compared with controls, whilst atypical oxidation of H80 and W32, as well as carboxymethyllysine (CML; K36) and glycation of K9, K23, K91, R115 and K122, was increased in this region. **g.** Distribution of residues exhibiting significantly higher levels of atypical modifications (oxidation, glycation). Residues in panels **e** and **g** are labelled using one letter amino acid codes with their side chains highlighted in red. Complete details of statistical analyses identifying PTM alterations are presented in **Supplementary Table 4**

Given the importance of copper binding for SOD1 structure [14], we next measured the isoelectric point (pI) of enzymatically-active SOD1 protein in the Parkinson disease SNc to evaluate the impact of decreased copper binding on SOD1 protein conformation. Changes in pI reflect altered amino acid solvent exposure, which can influence the aggregation propensity of SOD1, as well as the electrostatic guidance of superoxide towards the protein’s active site. Decreased SOD1 copper content was correlated with lower SOD1 protein pI in the SNc of Parkinson disease patients but not age-matched controls (**Fig. 1b**), indicating that conformational changes in SOD1 protein structure resulting from altered metallation are proportional in severity to the magnitude of copper deficiency in Parkinson disease. Collectively, these data suggest that decreased SOD1 copper binding may contribute to the accumulation of disSOD1 in the Parkinson disease SNc.

In addition to metal binding, the chemical modification of key amino acid residue side chains within SOD1 regulates structure (**Fig. 1c**), maturation, metallation, subcellular localization and antioxidant function of the protein under physiological conditions [14, 20]. Recognizing that disSOD1 pathology may result from either the disruption of these modifications or the introduction of atypical PTMs, we employed proteomic mass spectrometry to profile the post-translational fingerprint of SOD1 protein immunoprecipitated from the post-mortem SNc and OCx of Parkinson disease patients and age-matched controls [18]. While the SNc exhibits significant neuron death and disSOD1 pathology in Parkinson disease, neither of these features are abundant in the OCx [11], hence the inclusion of this brain region in PTM analyses enables the distinction of specific changes to SOD1 that may contribute to these factors in vulnerable brain regions. We identified 81 individual PTMs to 43 residues (28%) of SOD1 protein (154 amino acids) across both brain regions and diagnostic groups (**Fig. 1d; Supplementary Table 4)**, which were classified broadly as either of physiological significance (56 PTMs; acetylation, succinylation, phosphorylation, deamidation, ubiquitylation, glycosylation, acetylglucosamination) or being atypical modifications (25 PTMs; oxidation, nitration, glycation). These classifications were made based on their abundance in the healthy mammalian brain, with physiological PTMs being in high abundance and pathological PTMs being in low abundance, as well as their reported impact on SOD1 protein structure and function *in vitro* or *in vivo* [14]. Among physiological SOD1 PTMs, we observed a striking decrease in SOD1 O- and N-glycosylation and acetylglucosamination across 4 serine and asparagine residues (1.6-to-79.1-fold decrease) in the SNc of Parkinson disease patients compared with controls (**Fig. 1e,f; Supplementary Table 4)**, which would be expected to negatively impact nascent SOD1 protein folding and trafficking in this region [21]. By contrast, significant increases (1.9-to-6.8-fold) in 8 atypical SOD1 PTMs were identified in the SNc of Parkinson disease patients compared with controls (**Fig. 1f,g; Supplementary Table 4)**, all of which have been shown to induce structural disorder and aggregation of the protein *in vitro* [14, 22, 23]. Importantly, none of these atypical alterations was identified in the non-degenerating Parkinson disease OCx, suggesting they are specific to degenerating brain regions in Parkinson disease (**Supplementary Table 4)**. On the contrary, SOD1 glycosylation (13 residues; 1.9-to-6.7-fold increase) and deamidation (5 residues; 1.6-to-2.9-fold elevation) were upregulated in the OCx of Parkinson disease patients compared with controls (**Supplementary Fig. 1; Supplementary Table 5)**, which likely act to suppress the accumulation of disSOD1 in this brain region by improving protein folding, trafficking and turnover [24, 25]. It is unlikely that increased deamidation in this brain region induces structural disorder in SOD1 protein, as reported elsewhere [26], given we previously identified minimal disSOD1 in the Parkinson disease OCx [11].

Overall, we have identified a range of atypical post-translational modifications likely to impact the structural and functional integrity of SOD1 within the degenerating SNc in postmortem Parkinson disease brain, which have the capacity to underlie the selective accumulation of disSOD1 pathology in this region.

### A functional model of wild-type SOD1 pathology in the Parkinson disease brain

Cellular models cannot fully replicate the environment of the aged brain and neither misfolded SOD1, nor dopamine neuron death, can be quantified in the living human brain. Further, while transgenic SOD1 mice expressing mutant forms of the protein have been developed, they cannot be applied to study SOD1 pathology in Parkinson disease, which occurs in the absence of *SOD1* gene mutations [12]. To further study the relationship between wild-type disSOD1 pathology and nigral dopamine neuron health and function, we engineered a novel mouse strain expressing biochemical changes observed in the Parkinson disease SNc that we believe underlie the development of this pathology; decreased CNS copper levels and increased wild-type SOD1 protein expression [27]. Nigral copper deficiency in Parkinson disease patients is associated with a down-regulation of the cellular copper import protein Ctr1 [16], hence our novel mouse model was created by crossbreeding mice expressing half the normal level of Ctr1 (termed *Ctr1^+/-^* mice), with transgenic mice overexpressing human wild-type SOD1 (termed h*SOD1^WT^* mice) (**Fig. 2a**). The resulting novel strain of **<u>S</u>**OD1-**<u>O</u>**verexpressing **<u>C</u>**tr1-**<u>K</u>**nockdown mice will hereon be referred to as SOCK mice.

**Fig. 2.**
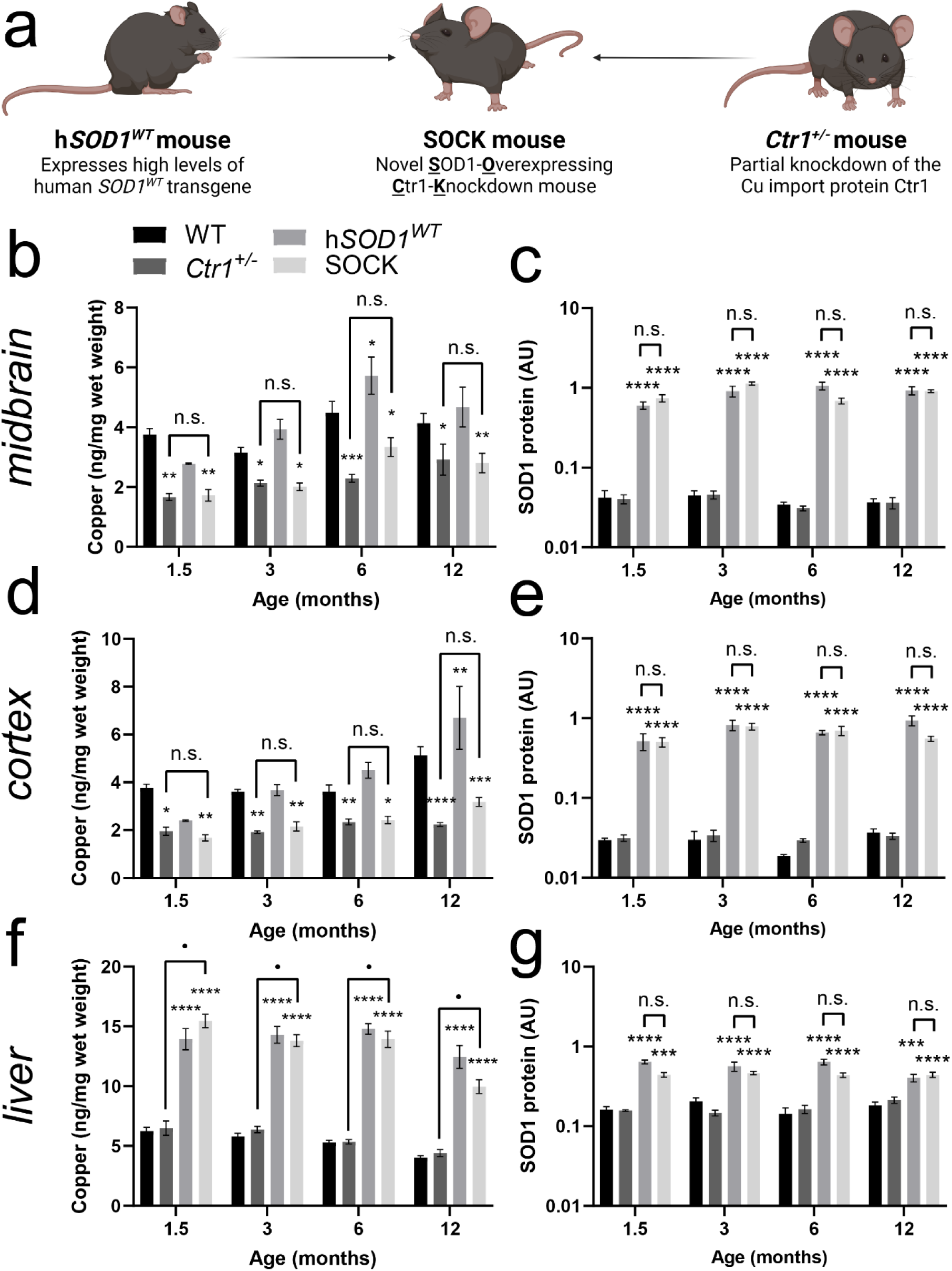
Novel SOCK mice recapitulate elevated SOD1 protein levels and brain copper deficiency observed in the post-mortem Parkinson disease SNc. **a.** Novel SOCK mice were developed by crossbreeding h*SOD1^WT^* mice, which overexpress wild-type human SOD1, with *Ctr1^+/-^* mice exhibiting decreased cellular copper within the central nervous system due to a knockdown of the neuronal copper import protein Ctr1. **b.** Midbrain copper levels quantified using inductively coupled plasma-mass spectrometry varied significantly between wild-type, h*SOD1^WT^*, *Ctr1^+/-^* and SOCK mice across all ages (two-way ANOVA: age – F_(3, 152)_ = 13.42, *p* < 0.0001; genotype – F_(3, 152)_ = 27.54, *p* < 0.0001), with decreases observed in *Ctr1^+/-^* and SOCK mice compared with wild-type mice (Dunnett’s multiple comparisons post hoc test: *Ctr1^+/−^* - 1.5 month, *p* = 0.0037, 3 month, *p* = 0.048; 6 month, *p* < 0.0001; 12 month, *p* = 0.032; SOCK - 1.5 month, *p* = 0.0065, 3 month, *p* = 0.023; 6 month, *p* = 0.022; 12 month, *p* = 0.0068). **c.** Levels of SOD1 protein in the midbrain quantified using immunoblotting varied significantly between all mouse strains (two-way ANOVA: F_(3, 119)_ = 748.6, *p* < 0.0001), with increases observed in h*SOD1^WT^* and SOCK mice compared with wild-type mice across all ages (Dunnett’s multiple comparisons post hoc test: *p* < 0.0001 for all comparisons). No differences were observed between h*SOD1^WT^* and SOCK mice at any age (Dunnett’s multiple comparisons post hoc test: *p* > 0.05 for all comparisons). Similar trends were observed for copper levels (**d;** two-way ANOVA: age – F_(3, 143)_ = 14.81, *p* < 0.0001; genotype – F_(3, 143)_ = 30.4, *p* < 0.0001. Dunnett’s multiple comparisons post hoc test: *Ctr1^+/−^* - 1.5 month, *p* = 0.02, 3 month, *p* = 0.0016; 6 month, *p* = 0.0097; 12 month, *p* < 0.0001; SOCK - 1.5 month, *p* = 0.0075, 3 month, *p* = 0.0051; 6 month, *p* = 0.028; 12 month, *p* = 0.0001) and SOD1 protein levels (**e;** two-way ANOVA: F_(3, 126)_ = 564.4, *p* < 0.0001. Dunnett’s multiple comparisons post hoc test: *p* < 0.0001 for all comparisons between h*SOD1^WT^* or SOCK mice and wild-type mice) in the cortex. **f.** Copper levels in the liver varied significantly between mouse strains (two-way ANOVA: F_(3, 150)_ = 361.8, *p* < 0.0001), with increases observed in h*SOD1^WT^* and SOCK mice compared with wild-type mice across all ages (Dunnett’s multiple comparisons post hoc test: *p* < 0.0001 for all comparisons). **g.** SOD1 protein levels in the liver varied significantly between mouse strains (two-way ANOVA: F_(3, 148)_ = 79.17, *p* < 0.0001), with increases observed in h*SOD1^WT^* and SOCK mice compared with wild-type mice across all ages (Dunnett’s multiple comparisons post hoc test: *p* < 0.0001 for all comparisons except 1.5-month-old SOCK mice (*p* = 0.0005) and 12-month-old h*SOD1^WT^* mice (*p* = 0.0003)). Data in panels **b-g** represent mean ± SEM. Sample sizes (/genotype/age group): 1.5 month = 5-8; 3 month = 7-13; 6 month = 10-16; 12 month = 6-14. Comparisons marked with an asterisk (*) denote comparisons made to wild-type mice, while those marked with a bullet point (•) demarcate those made between SOCK and *Ctr1^+/−^* mice. \**p* < 0.05, \*\**p* < 0.01, \*\*\**p* < 0.001, \*\*\*\**p* < 0.0001, • *p* < 0.0001, # *p* < 0.0001

Cellular copper homeostasis and SOD1 protein expression are tightly linked, with the expression of human SOD1 previously shown to promote elevated central nervous system copper levels in transgenic mutant SOD1 mice [28–30]. To confirm that h*SOD1^WT^* transgene expression did not modulate the severity of copper deficiency induced by *Ctr1^+/−^* knockdown in SOCK mice, we employed inductively coupled plasma–mass spectrometry (ICP-MS) and immunoblotting to quantify the levels of copper and SOD1 protein in all four mouse strains between 1.5 and 12 months-of-age. We observed a comparable 55% decrease in midbrain copper levels between 1.5 and 12 months-of-age in *Ctr1^+/−^* and SOCK mice compared with age-matched wild-type mice (**Fig. 2b**), consistent with previous data from *Ctr1^+/−^* mice [31, 32] and the Parkinson disease SNc [33]. Midbrain copper deficiency was accompanied by 22.1-fold higher levels of wild-type SOD1 protein on average between 1.5 and 12 months-of-age in SOCK mice compared with wild-type mice, which closely matched the magnitude of SOD1 protein overexpression in this region of the h*SOD1^WT^* mouse brain (22.7-fold, **Fig. 2c**; **Supplementary Fig. 2**), as well as in other transgenic mutant SOD1 mouse strains. Similar profiles for the levels of cellular copper (**Fig. 2d**) and SOD1 protein (**Fig. 2e**) were observed in the cortex of all four mouse strains mice.

Consistent with the presence of alternative copper import mechanisms in the periphery [34], copper content in the liver of h*SOD1^WT^* and SOCK mice was elevated 2.4-to-2.6-fold between 1.5 and 12 months-of-age compared with wild-type mice (**Fig. 2f**). This can likely be attributed to an increased demand for hepatic copper resulting from the 3-fold increase in SOD1 protein expression in this organ across all ages examined for both strains (**Fig. 2g**) [35]. Elevated SOD1 protein levels likely also underlie the significant increase in SOD1’s other metal cofactor, zinc, in all investigated regions of transgenic SOCK and h*SOD1^WT^* mice at all ages compared with non-transgenic wild-type or *Ctr1^+/−^* mice (**Supplementary Fig. 3**).

Collectively, these data indicate that h*SOD1^WT^* transgene expression does not modulate the severity of copper deficiency induced by *Ctr1^+/−^* knockdown in SOCK mice, confirming the successful induction of brain copper deficiency and wild-type SOD1 protein overexpression in these mice.

### Altered post-translational modification of SOD1 in SOCK mouse midbrain resembles Parkinson disease

Our data collected from post-mortem Parkinson disease tissues suggests that copper deficiency and increased SOD1 protein levels trigger the development of disSOD1 pathology by altering physiological SOD1 PTMs and promoting the addition of atypical PTMs to the protein. To assess whether similar PTM alterations were elicited by these factors in SOCK mice, we again employed proteomic mass spectrometry to profile atypical and physiological SOD1 PTMs in the midbrain of 6-month-old SOCK mice compared with the aged human SNc. Mirroring changes observed in the Parkinson disease SNc (**Fig. 1**), we identified significant increases in 10 atypical SOD1 PTMs in SOCK mice (**Fig. 3a**; **Supplementary Table 6**). Six of these changes were also present in the midbrain of h*SOD1^WT^* mice, suggesting they are associated with high levels of h*SOD1^WT^* transgene expression (**Supplementary Fig. 4, Supplementary Table 7**). The remaining 4 PTM alterations were unique to SOCK mice (**Fig. 3b**) and are associated with aggregation of the protein *in vitro* [22, 23], including oxidation of several metal binding histidine residues (H46, H48, H71; **Fig. 3c**) and glycation of a prominent solvent-accessible lysine residue (K23; **Fig. 3d**). In addition to higher levels of atypical SOD1 PTMs, we identified dysregulation of 51 physiological SOD1 PTMs in the SOCK mouse midbrain compared with the healthy aged human SNc (**Fig. 3e)**, 35 of which were also altered in h*SOD1^WT^* mice (**Supplementary Fig. 4**). The remaining 16 physiological PTM changes were unique to SOCK mice and were all down-regulated compared with SOD1 isolated from the healthy aged human SNc (**Fig. 3f,g)**, including a substantial decrease in SOD1 glycosylation mirroring those observed in the Parkinson disease SNc. These alterations signify substantial disruption of pathways governing SOD1 protein trafficking (glycosylation) [21] and turnover (deamidation, ubiquitylation), as well as maturation and catalytic activity (acetylation, succinylation, phosphorylation) [20], in the midbrain of SOCK mice, which may contribute to SOD1 protein dysfunction and misfolding [14, 20]. Collectively, PTM changes specific to SOCK mice largely recapitulate those observed in the Parkinson disease SNc (**Fig. 3h**).

**Fig. 3.**
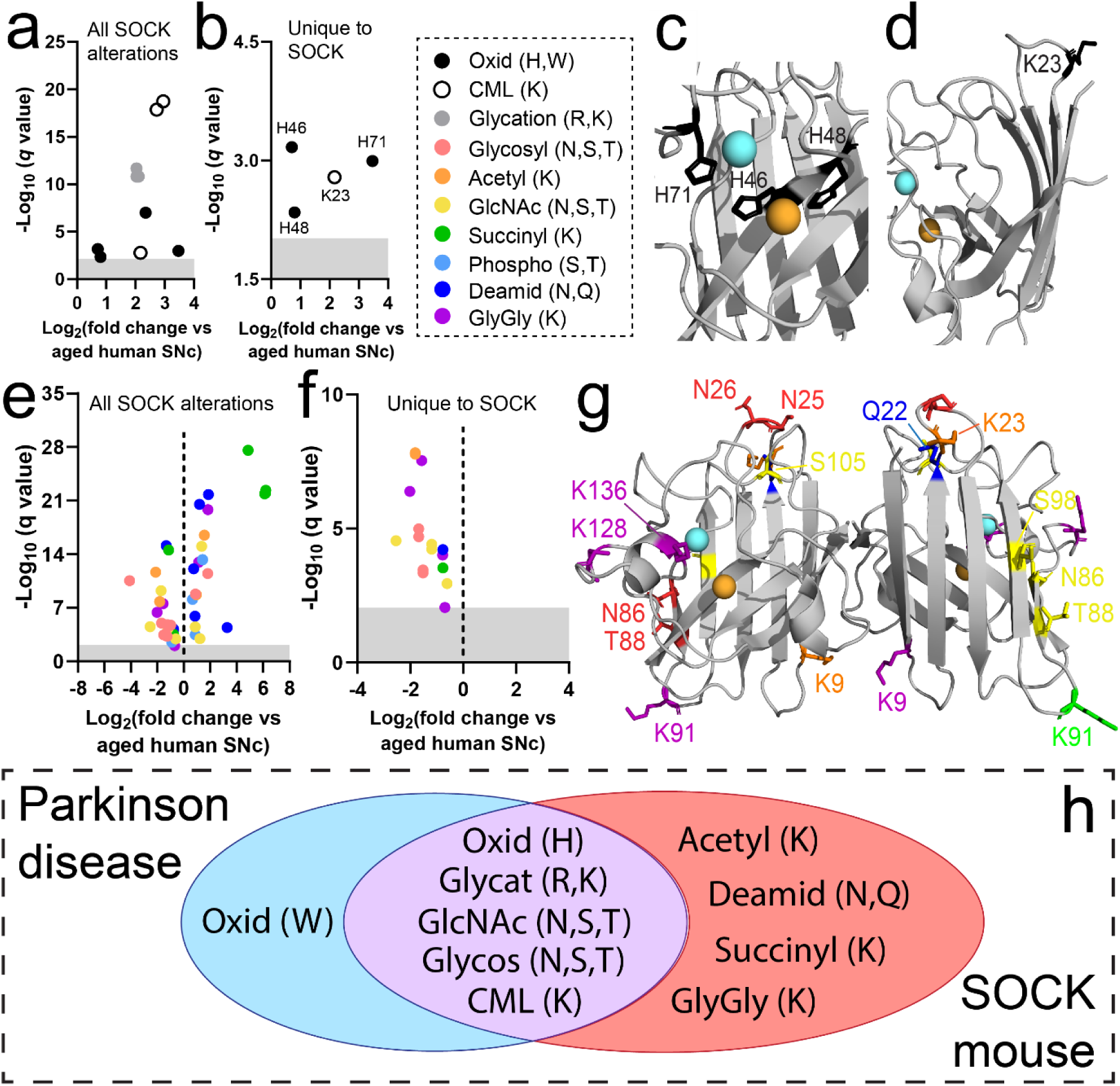
Post-translational modification of SOD1 is altered in the midbrain of SOCK mice. Atypical SOD1 PTMs were increased in the SOCK mouse midbrain (**a**), some of which were also shared with h*SOD1^WT^* mice, while others were unique to SOCK mice (**b**). These include oxidation of metal binding histidine residues (**c**; H46, H48, H71) and glycation of a solvent-accessible lysine residue (**d**; K23). Side chains of labelled residues are highlighted in black. Similar to atypical SOD1 PTMs, a number of physiological PTMs were significantly dysregulated in the SOCK mouse midbrain (**e**), some of which were shared with h*SOD1^WT^* mice, while others were unique to SOCK mice (**f**). GlyGly modifications result from tryptic digestion of ubiquitin-conjugated proteins, which serve as indicators of protein ubiquitination. **g.** Distribution of physiological SOD1 PTM alterations. **h.** SOD1 PTM alterations in the SOCK mouse midbrain largely overlap with those observed in the Parkinson disease SNc. Residues in panels **c, d** and **g** are labelled using one letter amino acid codes. Copper and zinc ions are highlighted in orange and cyan respectively. Complete details of statistical analyses identifying PTM alterations in SOCK and h*SOD1^WT^* mice are presented in **Supplementary Tables 6 and 7**. Abbreviations: CML, carboxymethllysine; Glycosyl, glycosylation; Acetyl, acetylation; GlcNAc, acetylglucosamination; Succinyl, succinylation; Phosphoryl, phosphorylation; Deamid, deamidation

### Enzymatic dysfunction and aggregation of wild-type SOD1 in the SOCK mouse SNc

Having established that SOD1 protein overexpression and copper deficiency promote SOD1 PTM alterations in SOCK mice matching those observed in Parkinson disease patients, we next sought to confirm that these changes result in enzymatic dysfunction and aggregation of the protein. Quantification of total SOD antioxidant activity in fresh frozen midbrain tissues revealed a 3.7-to-5-fold increase in SOD activity in h*SOD1^WT^* mice compared with wild-type mice across all ages (**Fig. 4a**), consistent with previous reports [36]. Total SOD activity was also increased in the SOCK mouse midbrain, albeit to a lesser extent (1.3-to-2.7-fold; **Fig. 4a**), suggesting cellular copper deficiency diminishes transgene-induced elevations in SOD activity. While total SOD activity includes the activity of all SOD enzymes (SOD2 and SOD3, as well as SOD1), we and others have demonstrated that SOD2 and SOD3 account for only 20% of total SOD activity in the healthy mammalian brain [11, 37]. This suggests that increased SOD activity in transgenic strains is likely derived almost exclusively from the increased levels of SOD1 protein expressed in these animals. The observed increases in SOD activity in the midbrain of both transgenic mouse strains were, however, disproportionately low considering the magnitude of SOD1 protein overexpression in this region. Indeed, normalization of total SOD activity measurements to SOD1 protein levels in the midbrain (**Fig. 2c**) revealed that SOD activity per unit of SOD1 protein was decreased by 76 - 85% in h*SOD1^WT^* mice compared with wild-type mice between 1.5 and 12 months-of-age. This deficit was significantly exacerbated by brain copper deficiency at all ages in SOCK mice, which exhibited an 89 - 93% decrease in SOD activity per unit of SOD1 protein compared with wild-type mice (**Fig. 4b**). Increases in SOD1 protein expression in h*SOD1^WT^* and SOCK mice were accompanied by increased levels of SOD1’s copper chaperone, CCS, (**Supplementary Fig. 5**), with a tight correlation observed between SOD1 and CCS protein levels in the SOCK mouse midbrain (**Supplementary Fig. 5**). We posit that the observed decrease in SOD activity per unit of SOD1 protein is therefore unlikely to derive from dysfunction of copper delivery mechanisms, but rather a mismatch between the demand for, and supply of, this essential cofactor. These data highlight an accumulation of immature, dysfunctional SOD1 in the midbrain of transgenic animals that is aggravated by copper deficiency.

**Fig. 4.**
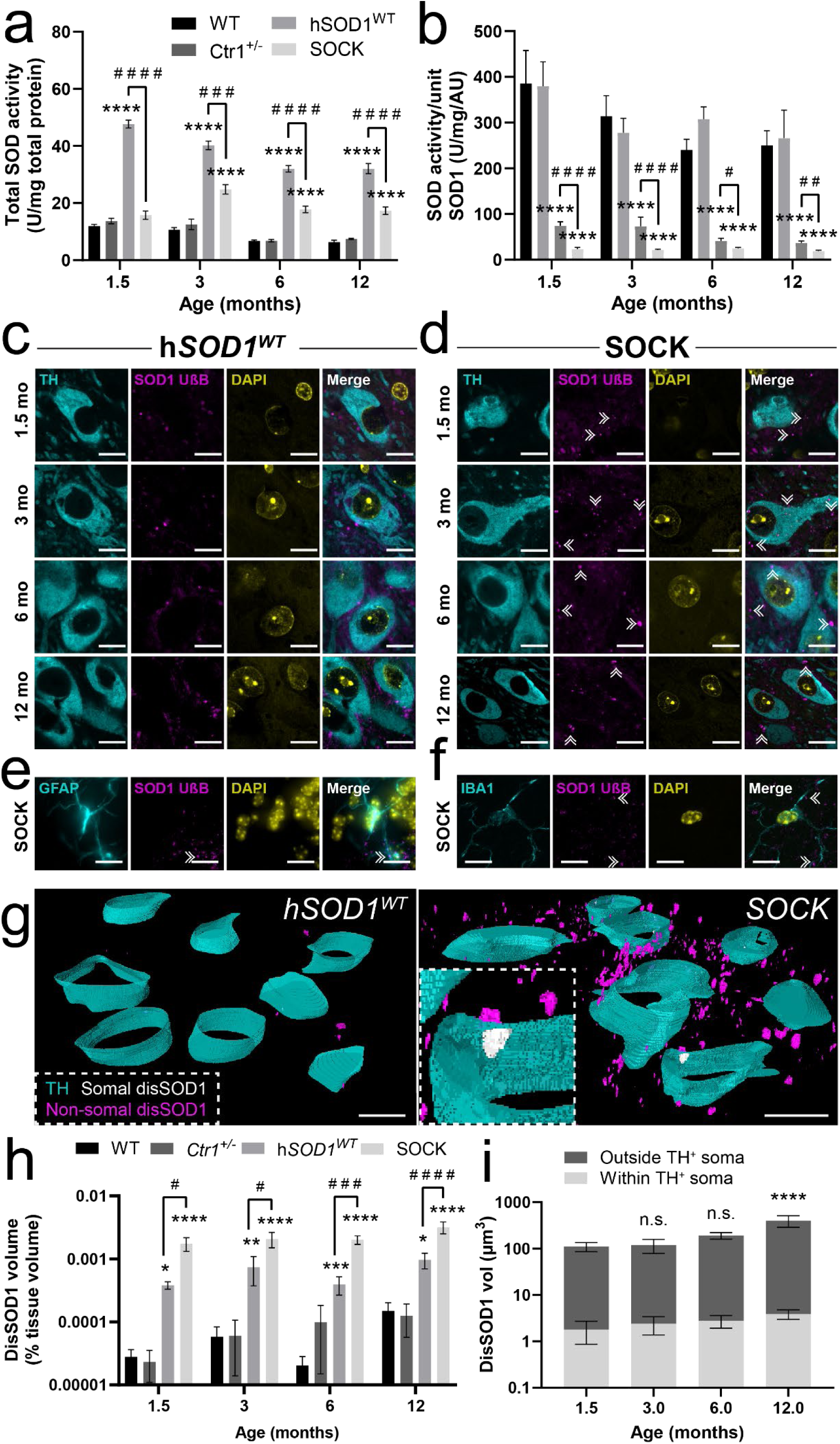
Immature, catalytically dysfunctional SOD1 accumulates and aggregates in the SOCK mouse SNc. **a.** Total SOD activity varied significantly in the midbrain between wild-type, h*SOD1^WT^*, *Ctr1*^+/−^ and SOCK mice aged 1.5-12 months old (two-way ANOVA: age – F_(3, 111)_ = 36.58, *p* < 0.0001; genotype – F_(3, 111)_ = 432.6, *p < 0*.0001). Activity was increased in the midbrains of both h*SOD1^WT^* and SOCK mice compared with wild-type mice at all ages except 1.5-month-old SOCK mice (Tukey’s multiple comparisons *post hoc* test: *p* < 0.0001 for all comparisons), yet was decreased in SOCK mice compared with h*SOD1^WT^* mice at all ages (Tukey’s multiple comparisons *post hoc* test: *p* < 0.0001 for all comparisons). **b.** SOD activity per unit of SOD1 protein also varied significantly in the midbrain between all four mouse strains (two-way ANOVA: F_(3,105)_ = 321.5, *p <* 0.0001) and was decreased in the midbrains of both h*SOD1^WT^* and SOCK mice compared with wild-type mice at all ages (Dunnett’s multiple comparisons *post hoc* test: *p* < 0.0001 for all comparisons) as well as in SOCK mice compared with h*SOD1^WT^* mice at all ages (Dunnett’s multiple comparisons *post hoc* test: 1.5 and 3 months, *p* < 0.0001; 6 month, *p* = 0.022; 12 month, *p* = 0.003). Immunofluorescent staining of fixed midbrain tissues from h*SOD1^WT^* (**c**) and SOCK (**d**) mice with the unfolded beta barrel (UβB) conformation-specific SOD1 antibody revealed the presence of disSOD1 aggregates (double white arrowheads) within and outside of dopamine neuron (tyrosine hydroxylase (TH)-positive) soma at all ages examined in SOCK mice, which was present at much lower levels in h*SOD1^WT^* mice. Corresponding images for *Ctr1^+/−^* and wild-type mice are presented in **Supplementary Fig. 6**. DisSOD1 staining was occasionally colocalized with the astrocyte marker, GFAP (**e**), but rarely with the microglial marker Iba1 (**f**). Images in panels **c - f** were acquired from 12-month mice. Antibody details are presented in **Supplementary Table 8**. **g.** Three-dimensional reconstruction of SOD1 aggregates in the SNc of 12-month-old h*SOD1^WT^* and SOCK mice (grey, TH-positive neuron soma; magenta, disSOD1 within TH-positive soma; yellow, extrasomal disSOD1). Scale bars represent 10 µm in panels **c-g**. **h.** The volume of disSOD1, expressed as a % of the volume of tissue within which it was quantified, varied significantly between genotypes across all ages (two-way ANOVA: age - F_(3,71)_ = 4.01, *p* = 0.011; genotype - F_(3,71)_ = 58.90, *p* < 0.0001) and was elevated in SOCK mice compared with h*SOD1^WT^* mice across all ages examined (Dunnett’s multiple comparisons *post hoc* test: 1.5 month, *p* = 0.039, 3 month, *p* = 0.019; 6 month, *p* = 0.0008; 12 month, *p* < 0.0001). **i.** Total disSOD1 volume varied within and outside of SNc dopamine (DA) neuron soma with age in SOCK mice (two-way ANOVA: age - F_(3,94)_ = 4.71, *p* = 0.0042; localization - F_(3,94)_ = 4.59, *p* = 0.0048) and was increased within DA neuron soma at 12 months-of-age compared with 1.5 months-of-age in these mice (*p* < 0.0001). Data in panels **a, b, h-i** represent mean ± SEM. Sample sizes (/genotype/age group): 1.5 month = 6-11; 3 month = 8-13; 6 month = 5-14; 12 month = 8-12. Comparisons marked with an asterisk (*) denote comparisons made to wild-type mice (or 1.5-month-old SOCK mice in panel **i**), while those marked with a hashtag (#) demarcate those made to h*SOD1^WT^* mice. # *p* < 0.05, ### *p* < 0.01, #### *p* < 0.0001, \**p* < 0.05, \*\**p* < 0.01, \*\*\**p* < 0.001, \*\*\*\**p* < 0.0001

A large body of data demonstrates that immature SOD1 species constitute key intermediates within pathways leading to SOD1 deposition [14, 38, 39], suggesting SOD1 aggregation accompanies alterations to SOD1 activity in the midbrain of SOCK mice. Using immunofluorescent staining of fixed brain tissue sections from the contralateral hemisphere of these same mice, we observed SOD1 puncta in the SNc (**Fig. 4c,d; Supplementary Fig. 6**) and cortex (**Supplementary Fig. 7**) of SOCK and h*SOD1^WT^* mice from 1.5 months-of-age onwards that were virtually absent in these regions of the *Ctr1^+/−^* and wild-type mouse brain at all experimental timepoints. Puncta were identified using well-characterized conformation-specific antibodies (**Supplementary Table 8**) that selectively recognize disSOD1 exhibiting an unfolded β-barrel (UβB; **Fig. 4c,d**) or exposed dimer interface (EDI; **Supplementary Fig. 8**), as well as pan-SOD1 immunostaining (**Supplementary Fig. 8**). Irrespective of detection antibody, disSOD1 puncta exhibited amorphous morphologies and ranged in size from 0.3–3 µm in diameter. These puncta were present within the soma of dopamine neurons in the SNc (**Fig. 4d**), as well as in astrocytes (**Fig. 4e**) and throughout the neuropil (**Fig. 4d**), but not microglia (**Fig. 4f**).

Three-dimensional reconstructions of these same midbrain tissue sections from z-stack images were generated to enable quantification of disSOD1 pathology within the SNc of all four mouse strains (**Fig. 4g; Supplementary Fig. 9**). These analyses revealed a significantly greater volume of punctate disSOD1 deposits in h*SOD1^WT^* and SOCK mice compared with wild-type and *Ctr1^+/−^* mice at all ages examined (**Fig. 4h)**, suggesting that higher SOD1 protein levels constitute a key contributor to disSOD1 proteinopathy. Furthermore, the volume of disSOD1 deposits was 2.8-to-5.1-fold greater in SOCK mice compared with h*SOD1^WT^* mice at all ages (**Fig. 4h)**, reinforcing the important role of copper deficiency in aggravating the accumulation of immature, dysfunctional SOD1 in the midbrains of SOCK mice. We posit that the absence of substantial disSOD1 aggregation in *Ctr1^+/−^* mice at any age (**Fig. 4h**) may indicate that copper deficiency alone does not trigger significant disSOD1 pathology, or alternatively suggests that mouse SOD1 may be comparatively resistant to demetallation-induced aggregation. This has not been investigated to date, although would perhaps derive from differences in the amino acid sequence of mouse SOD1 compared with the human isoform (84% sequence homology). Interestingly, incorporation of tyrosine hydroxylase-immunopositive cell bodies into these renders demonstrated that disSOD1 pathology was almost entirely (97.9-99%) localized outside of dopamine neuron soma in the SOCK mouse SNc at all ages (**Fig. 4i)**. These data suggest that SOD1 aggregation predominantly arises either within dopamine neuron processes, which could not be accurately incorporated into renders, in alternative non-dopaminergic cell types, or in the neuropil or extracellular space in the SNc of SOCK mice.

Overall, we provide clear empirical evidence that significant wild-type disSOD1 pathology accompanies altered SOD1 PTMs in an environment of copper deficiency and increased wild-type SOD1 protein overexpression, suggesting these pathways may drive disSOD1 pathology in the Parkinson disease SNc.

### SOCK mice exhibit age-dependent nigrostriatal degeneration in the absence of overt α-synuclein deposition

Having now established significant disSOD1 accumulation in the SNc of SOCK mice, which replicates similar pathology observed in the Parkinson disease SNc [11], we next sought to assess whether disSOD1 is associated with the two neuropathological hallmarks of Parkinson disease; nigrostriatal degeneration and synucleinopathy [40].

Stereological estimates of nigral dopamine neuron density in all four mouse strains were obtained using tyrosine hydroxylase immunostaining of serial brain tissue sections spanning the entire SNc. The raw neuron density for each mouse (**Supplementary Fig. 10**) was expressed as a percentage of the average density calculated for 1.5-month-old animals of that genotype (**Fig. 5a**) to enable better distinction of differences in the rate of nigral dopamine neuron loss with age between strains (**Fig. 5b**). While no significant differences were observed between h*SOD1^WT^*, *Ctr1^+/−^* and wild-type mice at any given age, SOCK mice exhibited a 23% loss of nigral dopamine neurons compared with wild-type mice at 6 months-of-age, which increased to 28% at 12 months-of-age (**Fig. 5a,b**). Relative to 1.5-month-old SOCK mice, these neuronal losses were as high as 49% by 12 months-of-age (**Supplementary Fig. 10**). No significant differences in SNc dopamine neuron loss were identified between SOCK and h*SOD1^WT^* mice until 12 months-of-age, at which point we observed a 21% greater loss of dopamine neurons within the SOCK mouse SNc compared with h*SOD1^WT^* mice (**Fig. 5a, Supplementary Fig. 10**). By applying linear regression to measured nigral dopamine neuron densities, we determined that wild-type and h*SOD1^WT^* mice would have exhibited 60% loss of these neurons by 21 months-of-age, compared to 17 and 14 months-of-age for *Ctr1^+/−^* and SOCK mice, respectively (**Supplementary Fig. 10**). These data highlight an important role for copper deficiency in accelerating age-dependent dopamine neuron degeneration in the mouse SNc, which is further augmented by SOD1 protein overexpression in SOCK mice. Delineation of dorsomedial, lateral and ventral nigral subregions revealed that the majority of dopamine neuron loss occurred within the dorsomedial SNc in SOCK mice (**Supplementary Fig. 11**), likely as this subregion occupies the greatest volume compared with other nigral subregions. Importantly, decreases in nigral dopamine neuron density were associated with higher disSOD1 pathological burden in SOCK mice (**Fig. 5c**), indicating disSOD1 pathology likely plays a key role in potentiating age-dependent nigral dopaminergic denervation in these mice.

**Fig. 5.**
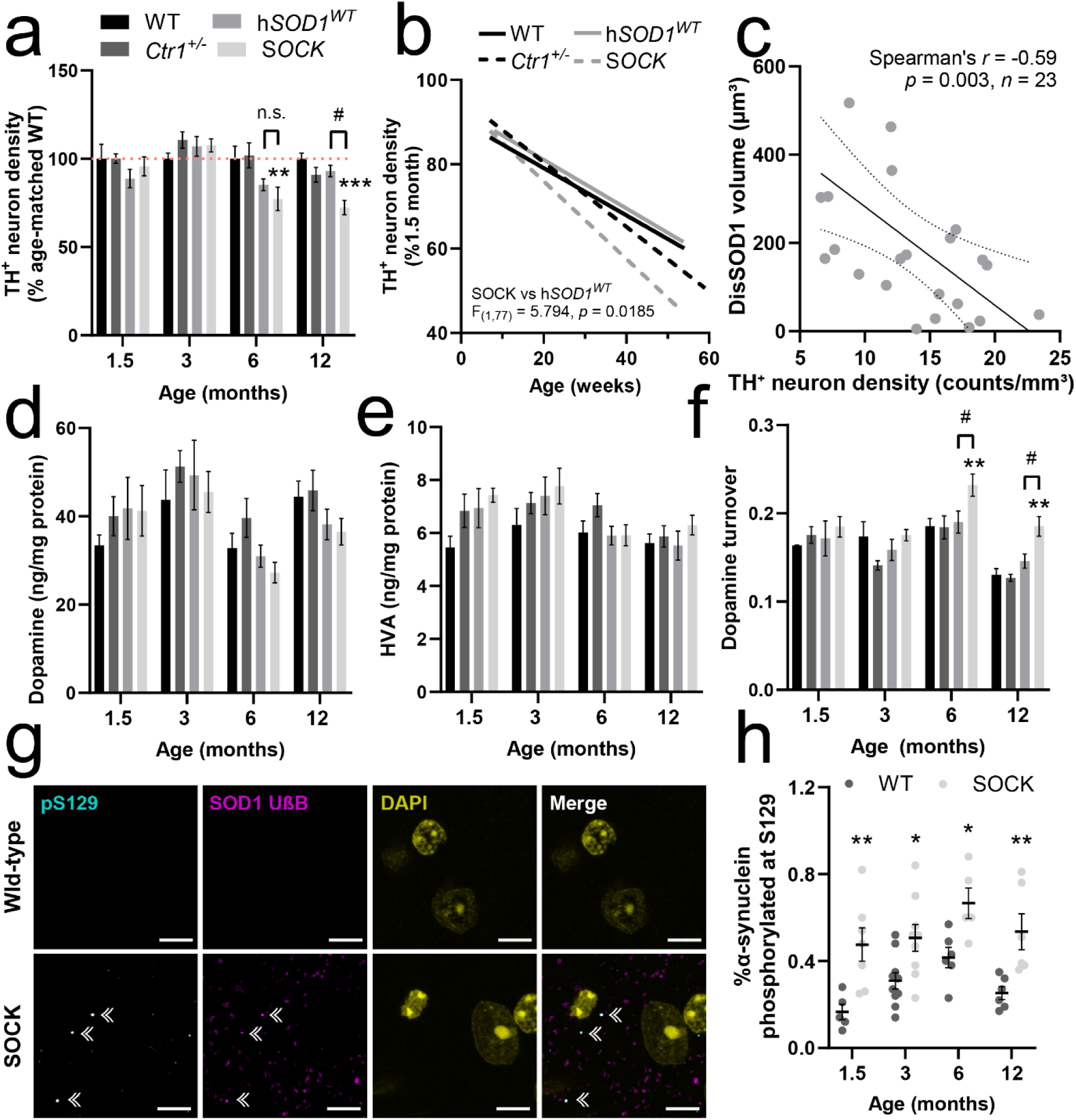
Dopamine neuron degeneration and perturbed dopamine metabolism occur in the absence of substantial α-synuclein aggregation in the SOCK mouse SNc. **a.** Quantitative stereology revealed significant variation in the density of dopamine neurons in the SNc of 1.5 – 12-month-old wild-type (WT), *Ctr1^+/^*, h*SOD1^WT^* and SOCK mice (two-way ANOVA: age – F_(3, 128)_ = 71.76, *p* < 0.0001; genotype – F_(3, 128)_ = 4.455, *p* = 0.0052), which was decreased in 6- and 12-month-old SOCK mice compared with WT mice (Dunnett’s multiple comparisons post hoc test: *p* = 0.0034 and 0.0002 respectively). The density of nigral dopamine neurons was also significantly lower in SOCK mice compared with h*SOD1^WT^* mice at 12 months-of-age (*p* = 0.021) but not 6 months-of-age (*p* = 0.70). Data represent the density of TH neurons as a proportion of the average number of TH neurons quantified for age-matched wild-type mice, with the red dotted line representing 100% of this neuronal density. Raw neuronal densities are reported in **Supplementary Fig. 10. b.** Nigral TH^+^ neuron density significantly decreased with age in all four mouse strains (linear regression, significantly non-zero; WT (F_(1,41)_ = 10.35, *p* = 0.0025), *Ctr1^+/−^* (F_(1,41)_ = 32.61, *p* < 0.0001), h*SOD1^WT^* (F_(1,22)_ = 8.86, *p* = 0.007) and SOCK (F_(1,36)_ = 34.45, *p* < 0.0001)), with a more rapid decrease observed in SOCK mice compared with h*SOD1^WT^* mice (statistics displayed in the panel). Data represent the density of TH neurons as a proportion of the average number of TH neurons quantified at 1.5 months-of-age for that same mouse genotype. **c.** Decreases in dopamine neuron density were correlated with higher disSOD1 pathological burden in SOCK mice (statistics displayed in panel). Striatal dopamine levels (**d**) were unchanged between genotypes across all ages (two-way ANOVA: F_(3, 148)_ = 1.444, *p* = 0.23), as were striatal levels of homovanillic acid (**e**, HVA; two-way ANOVA: F_(3, 145)_ = 1.914, *p* = 0.13). Striatal dopamine turnover (**f**; calculated by normalizing the amount of HVA to dopamine levels) varied significantly (two-way ANOVA: age - F_(3, 141)_ = 14.66, *p* < 0.0001; genotype - F_(3, 141)_ = 7.84, *p* < 0.0001) and was increased in 6- and 12-month-old SOCK mice compared with WT mice (Dunnett’s multiple comparisons post hoc test: *p* = 0.0073, 0.0007 respectively) and h*SOD1^WT^* mice (Dunnett’s multiple comparisons post hoc test: *p* = 0.034, 0.046 respectively). **g,** Immunofluorescent staining of fixed midbrain tissues from 12-month-old SOCK mice with antibodies recognizing pS129 α-synuclein and SOD1 in an unfolded beta barrel (UβB) conformation revealed a small population of aggregates containing both proteins in the SOCK mouse SNc (double white arrowheads) that was absent in the WT SNc. Scale bars represent 10 µm. Antibody details are presented in **Supplementary Table 8**. **h.** The proportion of α-synuclein phosphorylated at Ser129 was significantly increased in the SOCK mouse midbrain compared with WT mice across all ages (two-way ANOVA: age – F_(3, 46)_ = 4.08, *p* = 0.012; genotype – F_(1, 46)_ = 37.11, *p* < 0.0001; Sidak’s multiple comparisons post hoc test: *p* values for 1.5 – 12 months = 0.0045, 0.028, 0.038, 0.0098, respectively). Comparisons marked with an asterisk (*) denote comparisons made to wild-type mice, while those marked with a hashtag (#) demarcate those made to h*SOD1^WT^* mice. Data in **a, d, e, f, h** represent mean ± SEM, with *n* = 4–16/group. # *p* < 0.05, \**p* < 0.05, \*\**p* < 0.01

In addition to nigral dopamine neuron number, we assessed whether disSOD1 pathology impacted dopaminergic activity from the SNc by quantifying levels of dopamine and its metabolite, homovanillic acid (HVA), in the striatum of all mouse strains between 1.5 and 12 months-of-age using high performance liquid chromatography. While the total amounts of striatal dopamine and HVA were unchanged between mouse strains across all ages (**Fig. 5d,e**), we identified a 25% elevation in striatal dopamine turnover (HVA normalized to dopamine levels) in SOCK mice compared with wild-type mice at 6 months-of-age, which increased to 42% by 12 months-of-age (**Fig. 5f**). These rates of turnover were significantly higher than those observed in h*SOD1^WT^* and *Ctr1^+/−^* mice at 6 and 12 months-of-age, which were unchanged compared with wild-type mice. We posit that this indicates surviving nigral dopamine neurons in aged SOCK mice up-regulate dopamine synthesis to ensure consistent striatal dopamine release.

Hallmark Lewy pathology in Parkinson disease is largely comprised of aggregated α-synuclein protein phosphorylated at serine residue 129 (pS129) [41]. Immunostaining of fixed SOCK mouse midbrain tissues using an antibody specifically recognizing pS129 α-synuclein revealed a small proportion (< 4%) of disSOD1 aggregates also containing pS129 α-synuclein protein (**Fig. 5g**), consistent with a reported cross-seeding interaction between disSOD1 and pathological α-synuclein *in vitro* [42]. No pS129 α-synuclein aggregates lacking disSOD1 were identified. A 60-287% increase in the proportion of soluble pS129 α-synuclein in the SOCK mouse midbrain was also observed compared with wild-type mice across all ages (**Fig. 5h**), which may reflect an up-regulation of molecular pathways designed to lessen deposition and promote clearance of soluble aggregation-prone α-synuclein, previously demonstrated *in vitro* [43, 44].

Taken together, our data highlight an age-dependent relationship between disSOD1 pathology and nigrostriatal degeneration occurring in the absence of overt α-synuclein aggregation.

### Motor impairment in SOCK mice is not associated with spinal motor neuron degeneration

Age-dependent nigrostriatal degeneration is a major contributor to progressive motor impairment in Parkinson disease patients [40]. Given the identified changes in nigrostriatal function and structure in SOCK mice, we evaluated broad motor performance in our mice using the rotarod test. An age-dependent decline in motor function was observed in all strains, with SOCK mice performing consistently more poorly at all ages compared with wild-type mice (**Fig. 6a**), exhibiting an 18-57% faster time to fall on average between 1.5-to-12 months-of-age (**Fig. 6b**). Transgenic h*SOD1^WT^* mice also exhibited poor motor function, although this began at a later age than SOCK mice. Nonetheless, motor function in h*SOD1^WT^* mice declined at the same rate as SOCK mice until 12 months-of-age, when SOCK mice performed significantly poorer than h*SOD1^WT^* mice (**Fig. 6b**). No differences in rotarod performance were observed for *Ctr1^+/−^* mice compared with wild-type mice, consistent with recently reported data [32]. These data imply brain copper deficiency alone does not result in movement dysfunction, but may aggravate the effects of increased hSOD1^WT^ expression on motor function to trigger motor dysfunction earlier in life and accentuate deficits at older ages.

**Fig. 6.**
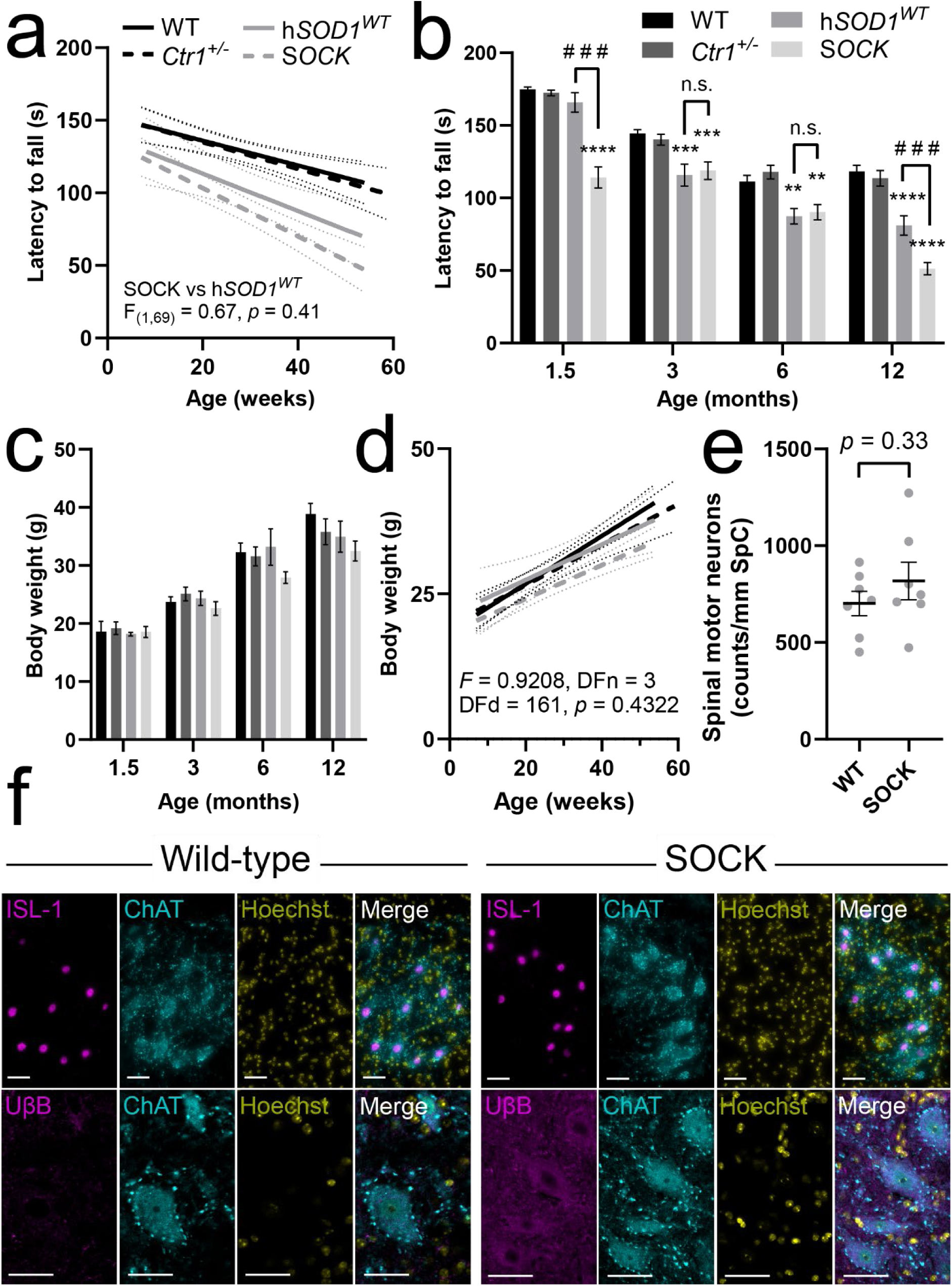
Neither body weight, nor spinal cord motor neuron density, are altered in SOCK mice despite poorer motor performance on the rotarod apparatus. Assessment of motor performance was conducted for wild-type (WT), *Ctr1^+/−^*, h*SOD1^WT^* and SOCK mice on an accelerating rotarod apparatus for a maximum of 180s. **a.** All mouse strains exhibited progressively poorer motor performance as they aged, with SOCK and h*SOD1^WT^* mice exhibiting a greater decline in performance compared with WT and *Ctr1^+/−^* mice. There was no difference in the rate of decline in motor performance between SOCK and h*SOD1^WT^* mice (statistics displayed in panel), resulting in significant differences between these strains and wild-type mice across most ages examined (**b;** two-way ANOVA: age - F_(3, 1008)_ = 89.91, *p* < 0.0001; genotype - F_(3, 1008)_ = 63.14, *p* < 0.0001; Dunnett’s multiple comparisons post hoc tests: *p* values for SOCK mice 1.5 – 12 months = <0.0001, 0.0001, 0.0021, <0.0001, respectively; *p* values for h*SOD1^WT^* mice 1.5 – 12 months = 0.82, 0.0004, 0.0012, <0.0001, respectively). SOCK mice also exhibited significantly poorer motor performance compared with h*SOD1^WT^* mice at 1.5 and 12 months-of-age (*p* values for 1.5 – 12 months = 0.0001, 0.98, 0.98, <0.0001, respectively). **c.** Body weight was measured as an index of general animal health throughout their lifespan (*n* = 3–16/group). No significant changes in body weight were observed between genotypes at any experimental age (two way ANOVA: genotype - F_(3, 153)_ = 2.23, *p* = 0.087), with all mouse strains gaining weight at a similar rate (**d**; statistical comparison of slopes presented in panel). **e.** The number of spinal motor neurons co-expressing choline acetyl-transferase (ChAT) and islet 1 (ISL-1) proteins did not differ between wild-type and SOCK mice at 6 months-of-age (unpaired *t*-test: *t* = 1.003, *df* = 12, *n* = 7/group, *p* = 0.33). **f.** Representative immunostaining of 6-month-old wild-type and SOCK spinal cord tissues for ChAT, ISL-1, and the unfolded β-barrel (UβB) conformation-specific disSOD1 antibody, counterstained with Hoechst. Scale bars represent 30µm. Comparisons marked with an asterisk (*) denote comparisons made to wild-type mice, while those marked with a hashtag (#) demarcate those made to h*SOD1^WT^* mice. Data in panels **b**, **c** and **e** represent mean ± SEM. \**p* < 0.05, \*\**p* < 0.01, \*\*\**p* < 0.001, \*\*\*\**p* < 0.0001, # # # *p* < 0.001

Robust motor impairment is a key feature of transgenic mutant SOD1 mice, which are commonly used to study inherited forms of ALS. These mice are characterized by aggressive weight loss following progressive spinal motor neuron degeneration and exhibit large SOD1-immunopositive inclusions within surviving spinal motor neurons [45]. To evaluate whether motor dysfunction in SOCK mice could be attributed to the development of an ALS-like phenotype, we began by tracking body weight in all four mouse strains between 1.5 and 12 months-of-age. A 100% survival rate was observed for all mouse strains and no significant differences in mouse weight were identified between strains at any experimental timepoint (**Fig. 6c**), with SOCK mice gaining weight at a comparable rate to WT mice (**Fig. 6d**). We next quantified the number of motor neurons in the anterior horn of the lumbar spinal cord in 6-month-old SOCK and wild-type mice using immunofluorescent co-staining for two markers of this neuronal population; choline acetyltransferase (ChAT) and Islet 1 (ISL-1) (**Fig. 6e,f**). The number of spinal motor neurons co-expressing ChAT and ISL-1 did not differ between SOCK and wild-type mice (**Fig. 6e**). Finally, we immunostained lumbar spinal cord tissue sections from these same animals using the UβB conformation-specific disSOD1 antibody to assess whether SOCK mice exhibit large SOD1-immunoreactive motor neuron inclusions. We identified an increase in diffuse disSOD1 immunoreactivity within spinal motor neurons, consistent with the accumulation of disSOD1 in the SOCK mouse brain, although did not identify any large disSOD1-immunoreactive inclusions within spinal motor neurons (**Fig. 6f**).

Overall these data highlight that spinal cord disSOD1 accumulation and broad motor deficits in SOCK mice occur in the absence of aggressive ALS-like weight loss, spinal motor neuron SOD1 inclusions and degeneration, and premature death observed in transgenic mutant SOD1 mice.

## Discussion

A disease-modifying treatment for Parkinson disease is urgently needed [46], with the exploration of new disease targets alongside longstanding factors such as α-synuclein likely to be of value [47]. We previously identified significant deposition of wild-type disSOD1 in the post-mortem Parkinson disease brain that, unlike widespread deposition of α-synuclein, was concentrated in degenerating brain regions [11]. These data, together with the well-documented role of abnormal SOD1 in neuronal death in rare genetic forms of ALS, suggest that disSOD1 pathology may constitute a valuable new disease target for slowing the progression of Parkinson disease. The development of such treatments will require two key scientific advances; an improved understanding of the molecular pathway(s) underlying this pathology in the Parkinson disease brain, together with the generation of an appropriate model to study the evolution and therapeutic modulation of this pathology within SNc dopamine neurons. In this study, we achieved both key advances by identifying several molecular pathways driving the development of wild-type SOD1 pathology, which we replicated to generate the first mouse model expressing this pathology [11, 16].

Our findings from the post-mortem Parkinson disease SNc suggest mismetallation and altered post-translational modification of key amino acid residue side chains constitute key mechanisms driving misfolding and aggregation of the wild-type protein in this disorder. Mismetallated SOD1 exhibits greater structural flexibility, which increases the solvent-exposure of residues normally buried within the protein and makes them more susceptible to atypical chemical modification [14, 20, 48]. This is in line with previous findings by our group demonstrating stoichiometric imbalance in Cu:Zn binding to SOD1 aggregates in the Parkinson disease SNc [19], reinforcing the role of copper deficiency in destabilizing protein structure and promoting the formation of unstable isoforms. We propose this underlies increased SOD1 oxidation and glycation in the Parkinson disease SNc in this study, both of which increase the propensity of the protein to aggregate [22, 23] and may contribute to the accumulation of disSOD1 pathology in this region (**Fig. 7**). Oxidation of histidine and tryptophan residues promotes non-amyloid oligomerization and aggregation of the protein [49, 50], whilst glycation of key lysine and arginine residues (including carboxymethyllysine) induces protein unfolding and loss of secondary structure [51] and is also associated with aggregation [52]. We posit such alterations may result from rampant oxidative stress in this region of the Parkinson disease brain, which may be further perpetuated by the misfolding of such an important cellular antioxidant protein (**Fig. 7**). Increases in atypical SOD1 PTMs that promote its aggregation may be compounded by dysregulation of SOD1 glycosylation in the Parkinson disease SNc, which likely signifies disruption of nascent, immature SOD1 trafficking [21]. Considering the majority of protein glycosylation occurs within the endoplasmic reticulum (ER) and Golgi apparatus to regulate protein trafficking and maturation [21], we speculate that disSOD1 accumulation in the Parkinson disease SNc may therefore be associated with ER stress and Golgi fragmentation reported in this brain region [53, 54].

**Fig. 7.**
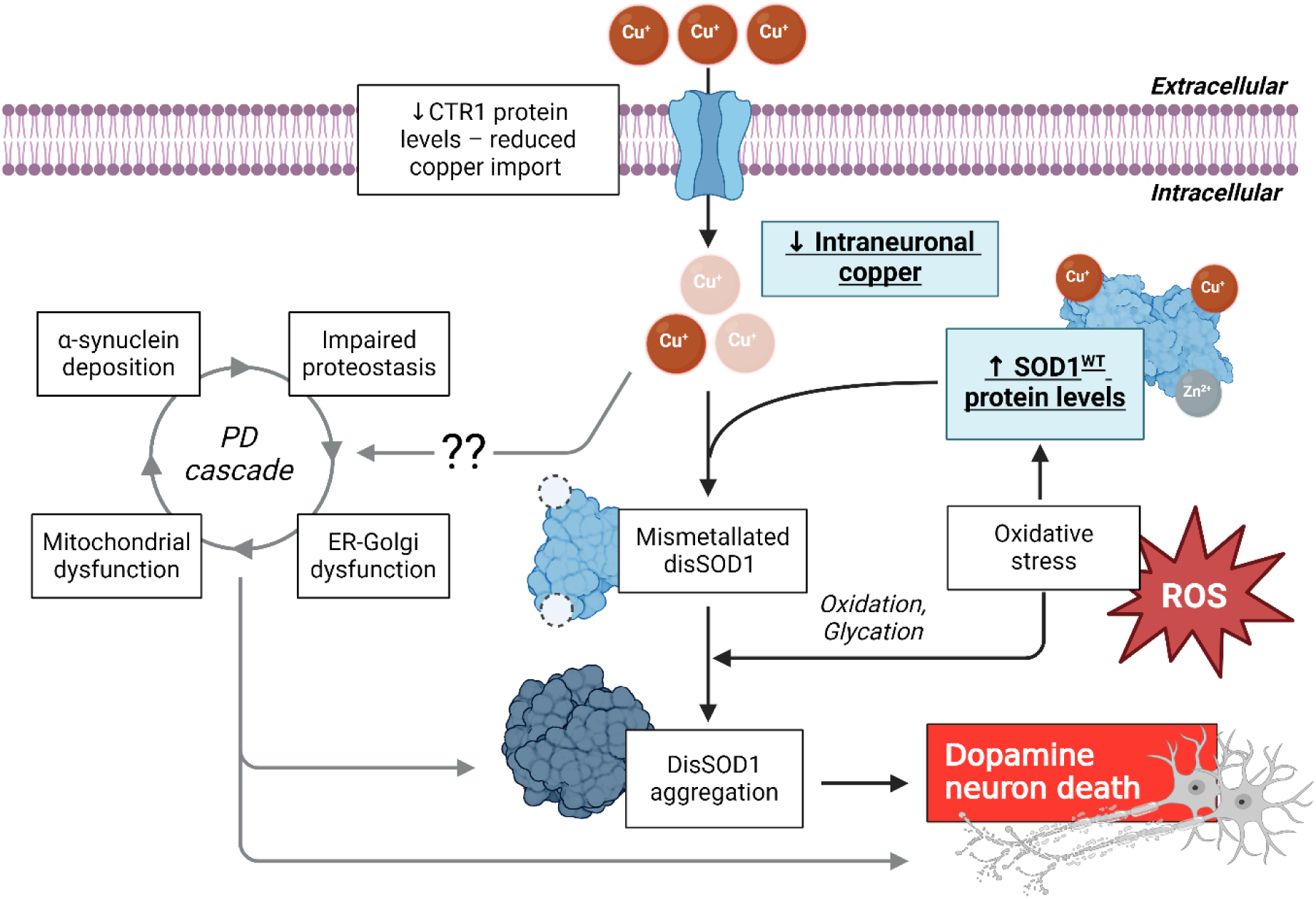
Proposed role of disSOD1 in nigral dopamine neuron death in Parkinson disease. Decreased expression of CTR1 in Parkinson SNc restricts neuronal copper import, limiting bioavailable copper and causing a decrease in copper binding to SOD1. This creates mismetallated disSOD1, which exhibits lower antioxidant activity per unit of protein and is prone to aggregation. Oxidative stress resulting from other etiological factors stimulates increased *SOD1* expression and promotes oxidation and glycation of solvent accessible residues within a now growing pool of disSOD1. Other pathologies may contribute to disSOD1 aggregation within the complex Parkinson disease (PD) degenerative cascade, which may themselves be exacerbated by brain copper deficiency. Combined, these changes may contribute to damage and death of dopamine neurons within the SNc.

The presence of similar alterations to SOD1 post-translational modifications in the midbrain of 6-month-old SOCK mice, alongside significant SOD1 misfolding, enzymatic dysfunction and deposition, reinforces the implication of these mechanisms in driving the development of wild-type disSOD1 pathology. This is especially likely given 6-month-old SOCK mice constitute a dynamic timepoint for disSOD1 accumulation, where significant pathology is already present but is still steadily increasing, whereas h*SOD1^WT^* mice possessing significantly less disSOD1 pathology lack these PTM changes. The presence of similar atypical oxidative modifications to SOD1, despite elevated total SOD activity, suggests that non-superoxide reactive oxygen species are responsible for oxidative damage to SOD1 in SOCK mice. Indeed, transgenic mutant SOD1 mice exhibit elevated hydrogen peroxide and hydroxyl radicals [55], with histidine residues particularly susceptible to oxidation by the latter of these species [56]. Paradoxically, this suggests that misfolded mutant and/or wild-type SOD1 may directly or indirectly induce redox imbalance to perpetuate their own misfolding.

Altered post-translational modification of SOD1 has long been suggested to constitute an important pathway driving mutant SOD1 deposition in rare inherited forms of ALS [48, 57], however our data provides the first *in vivo* evidence that copper deficiency and concomitant SOD1 protein overexpression promote the development of wild-type SOD1 pathology in the mammalian brain. The presence of a relatively low amount of this pathology within SNc dopamine neuron soma in 12-month-old SOCK mice suggests that disSOD1 deposition may initially develop away from the cell body in axons or dendrites, or may instead originate from alternative cell types such as astrocytes. DisSOD1 deposition was indeed observed in midbrain astrocytes in SOCK mice (**Fig. 3**), however it is unclear whether this reflects astrocytic uptake of disSOD1 from dying neurons, or alternatively whether this results from the migration of astrocyte-derived disSOD1 into neurons, which may induce their degeneration [58, 59]. In support of the latter of these pathways, selective expression of misfolded human mutant SOD1 in spinal motor neurons *in vitro* alone does not trigger motor neuron degeneration, however co-culture of these neurons with astrocytes expressing the same misfolded protein promotes their rapid degeneration [59]. Investigation of these pathways in future studies will be important to inform on mechanisms driving the development of wild-type disSOD1 pathology in SOCK mice, which may indicate tractable avenues for slowing or halting the development of this pathology.

Having identified potential mechanisms contributing to the development of wild-type disSOD1 pathology, we next explored the potential role of this pathology in Parkinson disease etiology. Our group previously identified a strong association between disSOD1 pathology and nigral dopamine cell death in Parkinson disease [11], which we reinforce in this study by demonstrating that disSOD1 pathology promotes significant nigrostriatal degeneration in our novel SOCK mouse model. We propose that the age-associated development of these features in SOCK mice demonstrates the value of these mice as a valuable alternative to more aggressive neurotoxin- or mutation-based mouse models of Parkinson disease, given this more closely resembles the progressive and age-dependent nature of nigrostriatal degeneration in idiopathic Parkinson disease patients [60]. Although the regional specificity of dopamine neuron death within the SNc of SOCK mice does not completely align with that observed in Parkinson disease patients, occurring largely in the dorsomedial SNc rather than the ventrolateral SNc [61], it is notable that genetic changes were engineered into these mice globally for the primary purpose of interrogating mechanisms resulting in disSOD1 pathology. Specific restriction of tissue copper deficiency and SOD1 protein overexpression to the SNc in future studies would better reflect the distribution of these changes in the Parkinson disease brain.

Importantly, the lack of significant body weight loss and spinal motor neuron degeneration in 12-month-old SOCK mice clearly distinguishes them from transgenic mutant SOD1 mouse models used to study ALS, which exhibit robust weight loss from 3 months-of-age following progressive spinal motor neuron degeneration and muscle wasting (∼5%/week [45]). This is particularly interesting given that mutant and wild-type SOD1 pathology in ALS and Parkinson disease exhibit common structural characteristics [11, 62], yet the global expression of human mutant SOD1 aggressively impacts spinal motor neurons, whilst the global induction of human wild-type SOD1 pathology results in nigral dopamine neuron death in the absence of spinal motor neuron degeneration. It must be acknowledged, however, that little-to-no attention has been paid towards nigrostriatal function in mutant SOD1 mice, hence we cannot discount the possibility that SNc dopamine neurons are also impacted by mutant SOD1 transgene expression, albeit likely to a much lesser extent. We posit that the ability of this pathology to underlie the degeneration of such distinct neuron populations may derive from the mechanisms of its formation. In SOCK mice, disSOD1 deposition was induced using biochemical alterations that characterize the Parkinson disease SNc, whilst similar pathology in ALS mouse models is induced by the introduction of known ALS-associated *SOD1* gene mutations.

In addition to nigrostriatal degeneration, previous studies have linked disSOD1 with the second pathological hallmark of Parkinson disease; α-synuclein deposition [42, 63]. These studies demonstrated a cross-seeding interaction between pathological forms of the two proteins in human cell lines [42] and mice [63], whereby mutant or aggregated α-synuclein promoted disSOD1 aggregation. However, these studies did not reciprocally examine the impact of disSOD1 on the subsequent development of α-synuclein pathology. We provide the first evidence that disSOD1 accumulation promotes higher α-synuclein S129 phosphorylation but not significant α-synuclein deposition, reinforcing findings which suggest that the co-deposition of these proteins within Lewy pathology in the Parkinson disease brain is more likely initiated by aggregated α-synuclein [42, 63].

The contribution of Parkinson-like disSOD1 pathology to progressive nigrostriatal degeneration identified in this study adds to mounting data suggesting that this pathology may constitute a novel therapeutic target for Parkinson disease. Several compounds that have been shown to combat SOD1-induced toxicity in transgenic mutant SOD1 mice are already underway in clinical trials for *SOD1*-linked ALS (ClinicalTrials.gov ID NCT04856982, NCT04082832), which we posit could represent potential means of combating disSOD1 pathology in Parkinson disease. However, while many approaches capable of mitigating mutant SOD1 pathology in *SOD1*-linked ALS rely on the complete ablation of SOD1 protein using gene silencing technologies [64], we propose that such approaches would be less effective in Parkinson disease where *SOD1* mutations are absent and oxidative stress is already rampant. We consider therapies capable of restoring the physiological maturation of SOD1, through improved metallation [65] or stabilization of the protein [66], to possess greater potential in mitigating nigrostriatal degeneration in Parkinson disease by simultaneously ameliorating SOD1 pathology and combating oxidative stress. Importantly, the low rate of success in translating potential disease-modifying therapies from animal models into the clinic across biomedical research is often attributed to differences between disease pathology in models and humans [67, 68]. The high degree of similarity in disSOD1 pathology between the SOCK mouse midbrain and Parkinson disease SNc highlights the suitability of this model as a drug screening tool for identifying therapies capable of slowing nigral dopamine neuron death by mitigating disSOD1 pathology.

### Conclusions

This study provides the first *in vivo* evidence that mismetallation and altered post-translational modifications drive the development of wild-type SOD1 misfolding, dysfunction and deposition. The recapitulation of these changes in SOCK mice using well documented biochemical features of the degenerating Parkinson disease SNc suggests that similar pathways may contribute to the development of this pathology in Parkinson disease patients. Furthermore, the direct association between disSOD1 pathology and age-dependent, progressive nigrostriatal degeneration in SOCK mice highlights a role for wild-type disSOD1 in the death of SNc dopamine neurons in Parkinson disease, advocating for the exploration of this pathology as a novel drug target for this disorder.

## Methods

### Human post-mortem tissues

Fresh frozen human post-mortem brain tissues from Parkinson disease patients (*n* = 19) and age-matched controls (*n* = 23) were obtained from the MRC London Neurodegenerative Diseases Brain Bank (6 Parkinson disease, 10 control; King’s College, London, UK), the Parkinson UK Brain Bank (4 Parkinson disease; Imperial College, London, UK) and the Sydney Brain Bank (9 Parkinson disease, 13 control; Sydney, Australia). Diagnoses of Parkinson disease were determined clinically by the donors’ physicians. Pathological identification of Lewy pathology and dopamine neuron loss in the SNc by brain bank neuropathologists confirmed clinical findings. All Parkinson disease cases were free of other neurological or neuropathological conditions. Genotyping confirmed the absence of *SOD1* mutations, as described below. Age-matched control cases were free of any clinically diagnosed neurological disorders and neuropathological abnormalities. Ethics approval was obtained from the University of Sydney Human Research Ethics Committee (approval number 2019/309). Demographic and clinical information for all cases are detailed in **Supplementary Table 1**. Diagnostic groups were matched for sex, age and post-mortem interval (**Supplementary Table 2**). Fresh frozen tissue samples were randomly numbered by a secondary investigator (V.C. or S.G.) prior to experimentation to blind primary investigators (B.G.T., A.A) to case diagnoses.

### SOD1 genotyping

*SOD1* genotyping was performed in 10 Parkinson disease and 10 age-matched control cases as previously described [69]. DNA was extracted from fresh frozen human brain tissue from the anterior cingulate or OCx using the DNeasy DNA extraction kit (Qiagen, Hilden, Germany, #69506), according to the manufacturer’s instructions. All five exons of SOD1, and at least 10 bp of flanking sequence were sequenced using PCR amplification and Sanger sequencing in duplicate. All results were independently analysed by two team members. Remaining Parkinson disease and control cases were genotyped previously [12].

### Animals

All methods conformed to the Australian Code of Practice for the Care and Use of Animals for Scientific Purposes [70], with protocols approved by the Animal Ethics Committee at the University of Melbourne (Ethics ID: 1814531.3) and ratified by the University of Sydney Animal Ethics Committees. Hemizygous male mice expressing seven-transgene copies of the human *SOD1^WT^* mouse strain (B6.Cg-Tg(SOD1)2Gur/J) were crossbred with female *Ctr1^+/−^* (*Slc31a1^tm2.1Djt^*/J) mice to produce the novel h*SOD1^WT^*/*Ctr1^+/−^* (SOCK) mouse strain. Both h*SOD1*^WT^ and *Ctr1^+/−^* mouse lines were sourced from The Jackson Laboratory (Bar Harbour, Maine, USA), and housed in filter top enclosures (12/12 h light/dark cycle, 22 °C, 45% humidity) with cardboard boxes and tubes for environmental enrichment. Enclosures contained Breeder’s Choice Cat Litter with paper tissues provided for bedding, and *ad libitum* access to standard chow pellets and water.

Tail snips were obtained from all mice prior to the age of weaning (3 weeks) for commercial genotyping of both *Slc31a1* and h*SOD1^WT^* genes. Mice were bred and aged to 1.5, 3, 6 and 12 months-of-age (group sizes displayed in **Supplementary Table 9**), before being anaesthetised with a lethal dose of xylazine (16 mg/kg body weight) and ketamine (120 mg/kg body weight). Mice were then perfused through the left ventricle with ice-cold 0.1 M phosphate buffer saline (PBS; pH 7.4, 4 °C) supplemented with phosphatase inhibitors (Phosphatase Inhibitor Cocktail 2; Sigma), protease inhibitors (Complete EDTA-free Tablets; Roche), and heparin (20 U/mL) and brain, lumbar spinal cord and liver tissues harvested. Brains were bisected sagittally into two hemispheres, before regions of interest (midbrain, striatum, cortex) were micro-dissected from the left hemisphere and stored with liver tissues at −80 °C for downstream biochemical analyses. The right brain hemisphere and entire lumbar spinal cord were placed in 4% paraformaldehyde overnight in preparation for immunohistochemical analyses.

### Fresh human and mouse tissue preparation for biochemical measurements

Fresh tissues were homogenized in 10x homogenization buffer volume (uL) per mg tissue weight (20 mM Tris-base pH 7.4 containing EDTA-free protease inhibitor (Sigma-Aldrich) and phosphatase inhibitor (Roche) using a Kontes pestle pellet mechanical tissue grinder (Sigma-Aldrich). Following homogenization, extracts were incubated at 4 °C for 30 min before protein concentration was determined using a bicinchoninic acid assay according to the manufacturer’s instructions (Thermo-Fisher Scientific).

### Measurement of enzymatically-active SOD1 metallation

The metal content of dimeric SOD1 was quantified using size-exclusion chromatography (SEC) coupled with native isoelectric focusing (nIEF) and synchrotron radiation X-ray fluorescence analyses, according to our published method [17, 18]. Briefly, brain tissue homogenates were subjected to SEC (Superdex 75 Increase 10/300 GL column connected to an Äkta Pure chromatography system, cooled to 4 °C, 100 mM ammonium acetate solution (pH 7.4) as the eluant) to purify enzymatically-active SOD1 dimers from fresh frozen post-mortem tissues. SOD1-containing fractions were then applied along immobilized pH gradient strips (ReadyStrip IPG Strips, pH 4-7, Bio-Rad) in duplicate and nIEF performed to further purify SOD1 according to its pI. SOD1 was identified within the first duplicate IPG strips using nitroblue tetrazolium (NBT) activity staining and metal quantification was performed on the second duplicate IPG strip at the pI of active SOD1. Copper and zinc quantification by SXRF was performed at the microprobe of the Hard X-ray Micro/Nano-Probe beamline P06 at the synchrotron PETRA III (DESY) in Hamburg (Germany)[71] using a Vortex silicon-drift X-ray detector with Cube preamplifier. SXRF analyses were carried out with an X-ray beam of 14 keV photon energy, 0.5×0.5 mm² beam size and 1.1×10^11^ ph/s and in-house code enabled online data analysis. Measurements were performed in triplicate or quintuplicate for each sample, which were averaged and used to generate mean Cu/Zn ratios and sd. Copper measurements represent total cuprous and cupric ions, while zinc measurements only represent Zn^2+^ ions as this is the only oxidation state of this metal. Limit of detection was calculated from blank measurements and only results above 3 LOD were retained for final analysis.

### SOD1 immunoprecipitation and preparation for proteomic mass spectrometry

SOD1 protein was immunoprecipitated from human and mouse brain tissue homogenates as previously described [18]. Briefly, 10 mg of Dynabeads M-280 Tosylactivated (Invitrogen, Carlsbad, CA, USA) were conjugated to 100 µg polyclonal SOD1 antibody (Enzo Life Sciences, Farmingdale, NY, USA; **Supplementary Table 8**) at 40 mg beads/mL in coupling buffer (0.1 M boric acid, pH 9.5; 1.2 M ammonium sulphate) overnight at 37 °C. Dynabeads were blocked with 0.5% BSA in PBS (pH 7.4), washed with 0.1% BSA in PBS (pH 7.4), and incubated with tissue homogenates (200 µg total protein) diluted in PBS (pH 7.4) to 40 mg beads/mL overnight at 4°C. Following PBS washes, immunoprecipitated proteins were eluted from Dynabeads using successive 10 min incubations with 0.1 M glycine (pH 3), eluants neutralized using an equivalent volume of ammonium bicarbonate (pH 8) and extracts dried under pressure using a vacuum concentrator. Dried SOD1 immunoprecipitates were resuspended in 50 mM ammonium bicarbonate (pH 8) containing 6 M urea, reduced with dithiothreitol (DTT; 10 mM final) for 30 min at 56 °C, alkylated with iodoacetamide (IAA; 20 mM final) for 30 min at room temperature in the dark, and finally quenched with a further 10 mM DTT for 30 min at room temperature. Samples were diluted five-fold using 50 mM ammonium bicarbonate (pH 8) to decrease the concentration of urea to 1.2 M and acetonitrile added (10% final), before in-solution digestion performed overnight at room temperature using 0.2 µg sequencing-grade modified trypsin (Promega, Madison, WI, USA). Samples were then acidified using trifluoroacetic acid, desalted using Pierce C18 Tips (ThermoFisher Scientific, USA) according to the manufacturer’s instructions, and dried under pressure using a vacuum concentrator. Samples were resuspended in loading buffer (0.1% formic acid, 3% ACN) and transferred to HPLC vials immediately prior to mass spectrometry analyses.

### Mass spectrometry data acquisition and analysis

Label-free Fourier Transform Mass Spectrometry was employed to analyse immunoprecipitated protein extracts at Sydney Mass Spectrometry (Sydney, New South Wales, Australia). Analyses were performed using an UltiMate 3000 RSLCnano system (ThermoFisher Scientific, USA) coupled online via a Nanospray Ion Source (ThermoFisher Scientific, USA) to a Q Exactive HF-X Hybrid Quadrupole-Orbitrap Mass Spectrometer (ThermoFisher Scientific, USA). Peptide digests were loaded onto an in-house packed ReproSil-Pur 120 C18-AQ analytical column (75µm id x 40 cm, 1.9 µm particle size; Dr Maisch, Ammerbuch, Germany) regulated to 60 °C using a PRSO-V2 Sonation column oven (Sonation, Baden-Wuerttemberg, Germany). A binary gradient of solvent A (0.1% formic acid in MilliQ water) to solvent B (0.1% formic acid in 80% ACN diluted with MilliQ water) was used for peptide elution at a separation flow rate of 300-450 nL/min over 90 min. The mass spectrometer operated in positive ion mode at a 2.4 kV needle voltage. Data were acquired using Xcalibur software (Version 4.4.16.14, ThermoFisher Scientific, USA) in a data-independent (DIA) mode. The MS was operated in a data-independent fashion with 20 dynamic DIA segments covering the mass range from 350 to 1650 *m/z*. The resolution for the MS1 scan was set to 60k with a max injection time of 50 ms and an AGC target of 3e6. The DIA scans were acquired in the Orbitrap with a resolution of 30k after fragmentation in the HCD cell (max injection time: auto; AGC target: 3e6; fixed first mass: 300 *m/z*; loop count: 1; MSX count: 1; isolation window: 26-589 *m/z*).

Raw DIA data was processed using Spectronaut software’s directDIA workflow (Version 19, Biognosys, Zurich, Switzerland), whereby raw data files were first used to generate project-specific spectral libraries *in silico* using Spectronaut Pulsar. Separate libraries were generated for each experimental question; the first used raw data files from Parkinson disease and control SNc extracts to address SOD1 PTM changes in Parkinson disease, while the second utilized raw data from SOCK and h*SOD1^WT^* mice, as well as human control SNc. Library generation was performed using BGS factory settings, with the Human Uniprot fasta file employed as a protein database for searches. Two missed cleavages were allowed and the false discovery rate (FDR) controlled at 1% for both PSM and protein group levels. Peptide identification and label-free quantification in each individual sample were then performed using default BGS factory settings, with spectra screened against Uniprot entry P00441 (SODC-HUMAN), corresponding to human SOD1. Cross-run normalization was disabled, PTM localization was enabled (summative PTM consolidation strategy and a probability cutoff of 0.75), and analyses were performed with a log2 ratio candidate filter of 0.58, a confidence (Q) candidate filter of 0.05 and multiple comparisons testing correction enabled. Carbamidomethyl (C) was included as a fixed modification, while modifications of interest were included individually in variable modifications alongside acetylation (N-term) and oxidation (M). PTMs of interest for comparison between Parkinson disease and control extracts were analysed in separate analysis batches, and included; acetylation (K; +42.01), acetylglucosamine (NST; 203.08) carboxymethyllysine (K; 58.01), deamidation (N/Q; +0.98), kynurenine (W; +3.99), glycation (K/R; +108.02 with neutral loss of three water molecules [72]), glycosylation (NST; +162.05), nitration (W; +44.99), oxidation (H/W; +15.99), phosphorylation (S/T; +79.97), succinylation (K; +100.02), ubiquitination (GlyGly, K; 114.04). A maximum of 5 modifications per peptide was allowed, with two missed trypsin cleavages. SOD1 protein was not identified in negative control immunoprecipitates prepared using Dynabeads that were not conjugated to our capture antibody, suggesting negligible false discovery of SOD1 protein in tissue extracts [18]. No differences in the relative levels of PTMs of interest were identified between immunoprecipitated and non-immunoprecipitated commercial SOD1 protein in a previous study [18], implying our immunoprecipitation protocol did not significantly alter PTMs of interest.

### SOD1 and CCS protein quantification

Immunoblotting for SOD1 and CCS proteins was performed by probing membranes first for CCS, then simultaneously probing membranes for SOD1 and GAPDH following stripping of CCS antibodies. Protein samples (0.5 μg for h*SOD1^WT^* SOD1 overexpressing and SOCK and 2.5 μg for Ctr1^+/−^ and WT mouse) were incubated in loading buffer (17.5% sodium dodecyl sulphate, 50% glycerol, 400 mM dithiothreitol (DTT), 0.3 M Tris-base (pH 6.8), 0.25% bromophenol blue) for 45 min at 56 °C to reduce and denature sample protein content, before being loaded onto 4–12% Bis–Tris Criterion pre-cast gels (Bio-Rad, Hercules, CA) and separated by sodium dodecyl sulphate–polyacrylamide gel electrophoresis in a Mini-PROTEAN Tetra Cell system at 180 V for 40 min at 4 °C (Bio-Rad). A region-specific loading control, prepared by combining equal amounts of protein from 18 samples of each region (3-month-old mice, n = 9 h*SOD1^WT^*-expressing, n = 9 WT for h*SOD1^WT^*), was loaded onto each gel. Separated proteins were transferred to Immobilon-PSQ PVDF (Millipore, Billerica, MA) membranes overnight at 9 V at 4 °C, before being air-dried overnight and then blocked in 5% skim milk (Bio-Rad, Hercules, CA) in phosphate buffer saline containing 0.1%Tween®20 (PBST)(Sigma-Aldrich, St. Louis, MO) for 1 hr at room temperature. Membranes were then incubated with primary antibody against CCS (1:2000, Rabbit anti-CCS, raised against peptides corresponding to aa 252-270 of the human CCS sequence, gifted by Dr Isil Keshin, Umeå University, Sweden) diluted in 1% skim milk in PBST overnight at 4 °C, followed by incubation with horseradish peroxidase (HRP)-conjugated goat anti-rabbit IgG (1:5000, Bio-Rad, Hercules, CA) for 2 hrs at room temperature. Protein signals were obtained using an ECL western blotting detection system (Bio-Rad, Hercules, CA) as per the manufacturer’s instructions, and developed using the iBright imaging system (Invitrogen). Membranes were then incubated in a stripping buffer (25 mM Glycine, 1.5% SDS, pH 2.0) to remove primary and secondary antibodies, re-blocked with PBST and 5% skim milk and incubated with primary antibody against rabbit anti-SOD1 (1:2000, Enzo, NY, USA)) and rabbit anti-GAPDH (1:10,000 Merck, G9545) overnight at 4 °C. Membranes were incubated with HRP goat anti-rabbit IgG (1:5000 Bio-Rad, Hercules, CA) and protein signals were obtained using an ECL western blotting detection system (Bio-Rad, Hercules, CA). Antibody details are presented in **Supplementary Table 8**. CCS, SOD1 and GAPDH signal intensities were quantified by densitometry using iBright analysis software v5.2.2 (Invitrogen). CCS and SOD1 values were first normalized to the corresponding GAPDH values and then normalized to the loading control value in each gel. GAPDH values were unchanged between genotypes at each age in all regions, validating the choice of this protein as a housekeeping gene (**Supplementary Fig. 12**).

### Quantification of tissue metal levels

Metal levels in soluble and insoluble tissue extracts were quantified using ICP-MS, according to our group’s published methods [73]. Twenty-to-thirty microliters of tissue homogenate were dried down and digested overnight using concentrated nitric acid (50 µL, 70%, Suprapur grade, Merk Millipore) at room temperature. Samples were then digested for a further 30 min at 70 °C, incubated with concentrated hydrogen peroxide (30%, VWH International, PA, USA) for 60 min at 70 °C, and then diluted to 2 mL with 1% nitric acid (1:10 v/v; Suprapur grade, Merk Millipore) prior to analysis. Total metal levels in each sample were measured in triplicate using a Perkin Elmer Nexion 300X Inductively Coupled Plasma Mass Spectrometer. Buffer controls containing 1% nitric acid were incorporated every 20 samples. Helium (4 mL/min) was used as a collision gas for the removal of polyatomic interferences. Measured mass-to-charge (*m/z*) ratios were 63 (Cu) and 66 (Zn). External calibration was performed using S24 multi-element standards (High Purity Standards, USA) diluted in 1% HNO3, while rhodium (Rh; m/z = 45) was used as reference element via online introduction with a Teflon T-piece. Measurements were background corrected to metal levels in buffer controls, adjusted for dilution factors and standardized against original wet tissue weights. Samples below the instrument’s limits of detection were excluded from analyses.

### SOD activity measurement

SOD1 antioxidant activity was quantified in tissue extracts using a commercial SOD Assay Kit (Cat. #19160, Sigma-Aldrich, USA) according to the manufacturer’s instructions [11]. Briefly, samples containing 2 µg protein were diluted serially between 10- and 1000-fold and the assay signal measured in triplicate. A bovine SOD standard was used to generate a standard curve relating SOD activity to assay signal, which was applied to sample dilution curves to obtain SOD activity measurements. Total SOD activity in each sample was normalized to SOD1 protein levels measured using immunoblotting, which yielded a measure of SOD activity per unit of SOD1 protein in each sample.

### Fixed mouse tissue preparation for immunostaining

Following overnight fixation in 4% paraformaldehyde, mouse brain and lumbar spinal cord tissues were incubated in 30% (w/v) sucrose solution in a 50 mL sample collection tube for 24–48 h until they sunk to the bottom of their container. They were then embedded in Tissue-Tek® optimal cutting temperature medium (Sakura Finetek, Nagano, Japan; #4583) and stored at −80 °C. Fifty micrometer thick serial brain tissue sections were then cut from Bregma 2.53 mm to 3.04 mm using an Epredia^TM^ CrytoStar^TM^ NX50 cryostat (ThermoFisher Scientific) to ensure the entire rostro-caudal SNc was collected, while the lumbar spinal cord was cut into serial 30 µm sections. These were divided into three free-floating section series and stored in 0.1 M PBS containing 0.02% sodium azide at 4°C until staining.

### Nigral dopamine neuron stereology

Free-floating brain tissue sections for dopamine neuron stereology were incubated for 30 min in citrate buffer (pH 6.0; Fronine) at 95 °C and were then cooled to room temperature before proceeding with 3,3-diaminobenzidine (DAB) staining. Endogenous peroxidase activity was quenched by pre-treating sections in a 3% H_2_O_2_ solution in 50% ethanol solution for 30 min at room temperature. Sections were washed in PBS-T, then blocked for 1 h at room temperature using 0.5% bovine serum albumin (BSA) and 1% casein in PBS-T. Sections were incubated in an anti-tyrosine hydroxylase (TH) primary antibody (Merck Millipore, USA, #AB152, 1:5000) at 4 °C. Primary antibodies were detected using a biotinylated goat anti-rabbit IgG secondary antibody (Vector Laboratories, USA; #BA-1000, 1:200), followed by a tertiary antibody (Vector Laboratories, USA; #PK-7200, 1:100) and visualised using DAB (Sigma-Aldrich, USA). Sections were washed and mounted onto Superfrost^®^ Plus silica glass microscope slides (ThermoFisher Scientific), and counterstained with cresyl violet prior to coverslipping. All images were captured using an Olympus VS120 Virtual Slide Microscope (Olympus, Japan) at 20× magnification with 4 µm depth intervals. Quantitative stereological analysis of TH-positive neurons in the SNc was completed using VS-DESKTOP (Olympus, version 2.91). Coronal plates of the anatomical distribution of the SNc and its lateral, dorsomedial, and ventral subregions were established according to published literature to ensure regions of interest were consistently defined in all imaged animals [74]. TH-positive neuron staining containing Nissl bodies in SNc subregions were counted. Neuronal density was calculated as the number of SNc dopamine neurons divided by the total volume of the SNc. Ten percent of images from each cohort were independently counted by two researchers to measure the interrater reliability of counts, demonstrating excellent interrater reliability of these measurements (Cronbach’s *α* = 0.953, *n* = 17).

### Immunofluorescent staining

Free-floating brain and spinal cord tissue sections for immunofluorescent staining were brought to room temperature and pre-treated with 0.3% Triton X-100 made in PBS (PBS-Tx) for 45 min to increase tissue permeability, before antigen retrieval performed using citrate buffer at 95 °C (Vector Laboratories, USA). After blocking with 10% normal horse serum made in 0.1% PBS-Tx, sections were incubated with appropriate primary antibodies (**Supplementary Table 8**) at 37 °C for 1.5 h and then at 4 °C for two nights. Following PBS-Tx washes, appropriate non-spectrally overlapping fluorescent secondary antibodies (**Supplementary Table 8**) were then incubated under the same conditions as the primary antibodies, before cell nuclei were stained with 4′,6-diamidino-2-phenylindole (DAPI) for 20 min at room temperature. Sections were mounted and coverslipped using SlowFade^TM^ Glass Antifade Mountant (ThermoFisher Scientific, USA). Image acquisition was performed using a Nikon C2+ Confocal Microscope System and Nikon NIS-elements software (Nikon, v5.20.02). Large-scale format images were first acquired at 10× magnification to select three regions of interest from three sections spanning the rostrocaudal SNc. A 60× magnification oil immersion lens was used to image those regions of interest at 1024×1024 pixel resolution with 0.3µm Z-step through the entire tissue thickness. A negative control was run by substituting the primary antibody with normal serum where no positive immunolabelling was detected, thus validating the absence of non-specific fluorescent signals (**Supplementary Fig. 13**).

### Quantification of SOD1 in nigral dopamine neuron soma

Confocal microscope images of brain tissue sections stained using SOD1 UßB, TH, and DAPI, were converted from ND2 format to 3DTIFF using Fiji (NIH, v1.53J). Images were loaded on Avizo^TM^ 3D (2021.2, ThermoFisher Scientific) and a subvolume of the image was extracted for downstream analysis. Segmentation of TH^+^ neuron cell bodies was performed by manually tracing a region of interest on every second slice and interpolating the remaining slices based on the traced regions of interest. All slices were visually checked to ensure the region of interest was captured correctly. Slices interpolated inaccurately were corrected using the brush tool or repeated with an increased number of manually segmented regions of interest. A threshold intensity for SOD1 was assigned to analyzed images to identify nd quantify SOD1 aggregates. Variable illumination and image intensity between images required an individual threshold for each image. Threshold intensities for SOD1 were individually assigned for each captured region of interest. Separate images for the TH^+^ neuron cell bodies and SOD1 protein threshold were exported in a 3DTIFF format. A Python script was subsequently applied to all analyzed images to extract 3D measurements of SOD1 aggregate volume and its proportion inside and outside TH^+^ neuron cell bodies. This script is publicly-available on GitHub (owner: Richard Harwood, repository: image_analysis_SOD1).

### Lumbar spinal cord motor neuron stereology

Mouse lumbar spinal cord sections (50 sections/mouse on average) for motor neuron stereology were immunostained for the motor neuron markers choline acetyltransferase (ChAT) and Islet-1 (ISL-1) and imaged using an Olympus VS200 virtual slide scanner (Olympus, Japan) at 40x magnification using the maximum intensity projection function. Spinal cord sections were confirmed as being from the lumbar region using anatomical features identified in an atlas of the mouse spinal cord [75]. Once images were obtained, left and right grey matter horns that were ventral to the spinal canal were then converted to .tiff files separately using QuPath (v0.5.1) software. Ventral motor neurons that were considered positive for both ISl-1 and ChAT were then segmented for counting by manual and automated methods using FIJI imaging software. Briefly, automated segmentation was performed via a FIJI software script that imported the multiplex images into FIJI and split them into single channel images. Nuclei that were immunopositive for ISL-1 (488nm channel) were then pre-processed using enhanced contrast (saturation: 0.2%, normalize histogram) and background subtraction (rolling ball radius:100 pixels), before image thresholding applied using the yen method, which was chosen as the most appropriate of 17 methods trialed. Fill holes and erode were applied to binary masks followed by segmentation using analyze particle analysis (size: 0.5-infinity, circularity: 0.1-1). To confirm double staining of motor neurons, segmented ISL-1 nuclei were then overlayed on ChAT immunopositive staining (647nm channel, min and max intensity: 50-700) and each ISL-1 nuclei particle expanded proportional to its respective feret diameter. Average 488nm and 647nm intensities and area measures were recorded in overlap zones, while background 647nm intensity was measured as a 500x 500-pixel box overlayed over areas of background on a subset of the data (30% images). Positive motor neurons were counted when they had both ISL-1 nuceli segmentation and proximal ChAT staining above background. Counts were then normalized to a 1mm length of spinal cord by dividing the counts by the combined length of spinal cord comprised by the sections counted (e.g. 50 sections of 30µm thickness = 1.5mm length of spinal cord). Manual segmentation of ventral spinal cord motor neurons was performed on 10% of the dataset to confirm reliability of the automated data produced, whereby motor neurons that were positive for both ISL-1 and ChAT staining were counted in the extracted images. Manual counting demonstrated excellent reliability for the automated segmentation (Cronbach’s α= 0.985 n= 75).

### Quantification of striatal dopamine levels and turnover

Fresh-frozen striatal mouse tissue was homogenized by pulse sonication (30% duty cycle, output 2, 2 × 30 second intervals) in a solution of 150 mM phosphoric acid and 500 µM diethylenetriaminepentaacetic acid. Total protein concentration of the tissue homogenate was assessed using the bicinchoninic acid assay according to manufacturer protocol (Thermo-Fisher Scientific). Homogenized tissue was centrifuged at 16,000 *g* for 45 min at 4 °C and the remaining supernatant was spun in 3 kDa cut-off Amicon® centrifugal filter tubes (Merck Millipore) at 14,000 *g* for 90 min at 4 °C. Filtrates were collected and stored at −80 °C for downstream processing.

Quantification of dopamine, 3,4-dihydroxyphenylacetic acid (DOPAC), and homovanillic acid (HVA) levels were conducted using reverse-phase high performance liquid chromatography system (HPLC), consisted of a pump module (Shimadzu Prominence LC-20AD, Shimadzu Corporation) coupled to a reversed phase Gemini C18 110 Å column (5 μm pore size, 150 × 4.6 mm; Phenomenex) and an electrochemical detector (Antec Leyden) with a glassy carbon-working electrode maintained at +0.82 V against a Ag/AgCl reference electrode. Both detector and column were maintained at 40 °C. Twenty microlitres of prepared samples were injected via a Prominence autosampler (Shimadzu Prominence SIL-20A, Shimadzu Corporation, Kyoto, Japan). The mobile phase consisted of 0.01 M sodium phosphate monobasic (84% (v/v)) and methanol (16% (v/v)) containing 0.1 mM ethylenediaminetetraacetic acid, 0.65 mM 1-octanesulfonic acid and 0.5 mM triethylamine. Mobile phase was adjusted to pH 3.8 using 1 M hydrochloric acid and filtered through 0.22 μm Whatman® filter circles then pumped at a flow rate of 1 mL/min. Calibration curves prepared from starting concentrations of analytical standards of dopamine (1 μM), DOPAC (0.5 μM) and HVA (3 μM) were run at the beginning of each day to quantify any variability in the HPLC system. The Shimadzu integrated workstation LabSolutions software (Version 5.57; Shimadzu) was used to calculate the area under the curve for each peak of interest, and data subsequently normalized to the total protein concentration of each sample. Dopamine turnover was calculated as the ratio of HVA to dopamine. Neurotransmitter concentrations are expressed as ng/mg of protein (mean ±SEM).

### Quantification of α-synuclein phosphorylation

Total and pS129 α-synuclein were quantified in mouse midbrain tissue extracts using AlphaLISA SureFire Ultra assays for these targets (Revvity, Total α-synuclein; ALSU-TASYN-B, Phospho-α-synuclein (Ser129); ALSU-PASYN-B) according to manufacturer’s instructions. We first identified optimal dilution factors required to bring total and pS129 α-synuclein levels in extracts to within the linear dynamic range of the total and pS129 assay (25-fold for pS129 assay, 125-fold for total assay). Extracts were then diluted in assay lysis buffer to their optimal dilution factors and applied to total and pS129 α-synuclein assays. Total α-synuclein concentration in samples was calculated using a standard curve generated by measuring serial dilutions of purified wild-type α-synuclein (RP-009; Proteos, Kalamazoo, MI, USA) using the total α-synuclein kit. The concentration of pS129 α-synuclein in these samples was calculated using a standard curve generated by measuring serial dilutions of purified pS129 α-synuclein (RP-004; Proteos) using the pS129 α-synuclein kit. Concentrations in diluted samples were then multiplied by their dilution factors and expressed as ng/mg total protein to account for differences in protein concentration in the original extracts. The proportion of α-synuclein phosphorylated at S129 was calculated by dividing ng pS129 α-synuclein/mg total protein by the ng total α-synuclein/mg total protein, expressed as a percentage.

### Animal weight and locomotor function

At each age of interest, animal weight was recorded prior to the assessment of motor function using the rotarod test according to published methods [57]. Mice were initially habituated on the stationary rod, and then trained to walk on the rotating rod for 5 days prior to blinded data collection. Mice were tested on the rotarod twice a week during the data collection phase of the study. The rotation speed of the dowel was accelerated from 4 to 40 rpm over 180 sec and the length of time spent on the dowel (latency to fall) was recorded up to a maximum of 180 sec.

### Statistical analyses

Statistical analyses were performed using RStudio (Build 524) and SPSS 28.0 (IBM Corp, NY, USA). Outliers were defined by SPSS as ‘extreme values’ ≥3× the interquartile range (or ±2 standard deviations) and excluded from the analysis. Parametric tests or descriptive statistics with parametric assumptions (standard one- and two-way ANOVA, Pearson’s *r* and t-test) were used for variables meeting the associated assumptions, with data normality assessed using the Shapiro-Wilk test, Levene’s test, and Brown–Forsythe tests. One- and two-analysis of variance (ANOVA) were paired with Dunnett’s multiple comparisons post hoc test to assess pairwise comparisons between experimental groups for a given variable. Non-parametric tests or statistics (Kruskal–Wallis test and Mann–Whitney U-test) were used for variables where the observed data did not fit the assumptions of parametric tests, with Dunn’s multiple comparisons post hoc tests to assess pairwise comparisons between select diagnostic groups for a given variable. Where large differences between groups existed (e.g. SOD1 protein expression, SOD1 proteinopathy), data were log transformed prior to application of statistical tests. Cronbach’s alpha was used to measure interrater reliability between researchers performing quantitative stereology. Significance level was defined as p < 0.05 for all statistical tests. Graphs were generated using GraphPad Prism 9.4.0 (Graph-Pad, CA, USA).

## Abbreviations

ALS: amyotrophic lateral sclerosis
BSA: bovine serum albumin
CCS: copper chaperone for SOD1
CNS: central nervous system
Ct: control
Ctr1: copper transport protein 1
DIA: data independent acquisition
disSOD1: disordered SOD1
DOPAC: 3,4-dihydroxyphenylacetic acid
EDI: exposed dimer interface
ER: endoplasmic reticulum
HPLC: high performance liquid chromatography
h*SOD1^WT^*: wild-type human SOD1 (gene)
HVA: homovanillic acid
ICP-MS: inductively coupled plasma mass spectrometry
IPG: immobilized pH gradient
NBT: nitroblue tetrazolium
nIEF: native isoelectric focusing
OCx: occipital cortex
PD: Parkinson disease
pI: Isoelectric point
pS129: phosphorylated serine residue 129
PTM: post-translational modification
SEC: size-exclusion chromatography
SEM: standard error of the mean
SNc: substantia nigra pars compacta
SOCK: SOD1-overexpressing Ctr1-knockdown (mice)
SOD1: Cu-Zn superoxide dismutase 1
SOD2: Mn superoxide dismutase
SOD3: extracellular superoxide dismutase
SpC: spinal cord
SXRF: synchrotron X-ray fluorescence
TH: tyrosine hydroxylase
UβB: unfolded beta barrel
WT: wild-type.

## Declarations

### Ethics approval and consent to participate

All experimental procedures involving the use of human post-mortem tissues were approved by the University of Sydney Human Research Ethics Committee (approval number 2019/309). All experimental procedures involving the use of mice conformed to the Australian Code of Practice for the Care and Use of Animals for Scientific Purposes, with protocols approved by the Animal Ethics Committee at the University of Melbourne (Ethics ID: 1814531.3) and ratified by the University of Sydney Animal Ethics Committees.

## Consent for publication

All authors read and approved the final manuscript prior to publication.

### Availability of data and materials

All associated data from this manuscript are available from the corresponding author upon reasonable request. The script used for stereological quantification of nigral dopamine neurons is publicly-available on GitHub (owner: Richard Harwood, repository: image_analysis_SOD1; https://github.com/RichardHarwood/image_analysis_SOD1).

### Funding

This project was funded by the Michael J Fox Foundation for Parkinson Research in partnership with the Shake It Up Australia Foundation (MJFF-009200), as well as the National Health and Medical Research Council of Australia (APP1181864) and the University of Sydney. A.H.A was supported by Research Training Program Stipend Scholarship (SC1999) from the National Health and Medical Research Council. D.P.B. was supported by the Australian Research Council Discovery Project grant DP230101740. R.O. and S.R. were supported by the Association France Parkinson.

### Authors’ contributions

K.L.D. conceptualized, funded and supervised the study with assistance from A.A., B.G.T. and P.J.C. Animal ethics was obtained by P.J.C., while human ethics was obtained by B.G.T. and K.L.D. Human post-mortem tissues were obtained by B.G.T and SOD1 PTM data collected from these tissues by B.G.T., S.R., G.F., D.B. and R.O. Genotyping of SOD1 in human post-mortem tissues was performed by S.C.M.F., S.W., J.F. and I.B. Generation, maintenance, monitoring and behavioural analysis of all mice involved in this project was performed by S.N. with assistance from P.J.C., with tissue collection conducted by S.N. with assistance from P.J.C., K.K., J.R.L., K.L.D. and B.G.T. Mouse tissue processing for fixed and fresh tissue analyses was performed by B.G.T., B.R. and A.A. Collection of histological data from fixed mouse tissues was performed by A.A. with assistance from F.K., V.C., M.P., S.G., M.R., G.S. with analyses completed by A.A., R.H., V.C. and M.R. Collection of biochemical data from fresh frozen mouse tissues was performed by B.G.T., A.A., D.M., V.C., J.S., B.R., A.A.L., N.P., B.A., K.K., T.E.L. and D.P.B., with analyses completed by B.G.T and A.A. Manuscript writing was completed by A.A., B.G.T. and K.L.D. with input from all authors.

## Acknowledgements

Tissues were received from the Imperial College London Brain Bank, which is supported by Parkinson UK and the Multiple Sclerosis Society of Great Britain and Northern Ireland, as well as from The London Neurodegenerative Diseases Brain Bank, which receives funding from the Medical Research Council and the Brains for Dementia Research program, jointly funded by Alzheimer’s Research UK and Alzheimer’s Society. The authors acknowledge the facilities, as well as the scientific and technical assistance, of the Australian Microscopy and Microanalysis Research Facility (http://ammrf.org.au) node at the University of Sydney, as well as those of the Sydney Mass Spectrometry Facility at the University of Sydney. We acknowledge DESY (Hamburg, Germany), a member of the Helmholtz Association HGF, for the provision of experimental facilities. Parts of this research were carried out at PETRA III and we would like to thank Jan Garrevoet for assistance in using the P06 Microprobe. Beamtime was allocated for proposal I-20230761 EC. The authors would like to thank Bertrand Thomas, Yann Rufin and Karina Khambatta (University of Bordeaux, France) for their assistance in measuring SOD1 metallation, and Eunice Choi for her assistance in quantifying α-synuclein.

