## Supplementary Fig for "Impaired maturation of wild-type superoxide dismutase 1 associated with neurodegeneration in Parkinson disease brain and a novel murine model"

**SUPPLEMENTARY INFORMATION**

Supplementary Tables 1-9

1. Demographic and clinical information for human post-mortem tissue cases.
2. Demographic statistics for diagnostic groups.
3. Genetic screening of the *SOD1* gene in PD patients and controls.
4. SOD1 PTMs in post-mortem PD and control substantia nigra.
5. SOD1 PTMs in post-mortem PD and control occipital cortex.
6. Altered SOD1 PTMs in the SOCK mouse midbrain compared with healthy aged human SNc.
7. Altered SOD1 PTMs in the h$SOD1^{WT}$ mouse midbrain compared with healthy aged human SNc.
8. Primary and secondary antibody details for all methods used in this study.
9. Sample size of mouse groups in each genotype and age cohort.

Supplementary Fig.s 1-13

1. Altered SOD1 post-translational modifications in the post-mortem PD OCx.
2. Representative SOD1 and GAPDH immunoblots from midbrain tissues of all four mouse genotypes
3. Zinc levels in the midbrain, cortex and liver of all 4 mouse genotypes.
4. Altered SOD1 post-translational modifications in the h$SOD1^{WT}$ mouse midbrain.
5. Representative CCS immunoblots and quantification.
6. Midbrain disSOD1 immunostaining in 12-month WT and $Ctr1^{+/-}$ mice
7. Cortical disSOD1 pathology in 12-month SOCK mice.
8. Cross-validation of disSOD1 pathology detection in the SOCK mouse SNc.
9. Three dimensional reconstructions of disSOD1 aggregates in the SNc of all mouse strains.
10. Stereological estimates of SNc dopamine neuron density in all mouse genotypes.
11. Stereological estimates of dopamine neuron density in dorsomedial, lateral and ventral subregions of the SOCK mouse SNc.
12. GAPDH protein levels in the midbrain, cortex and liver of all mouse strains.
13. Negative control immunofluorescent staining.

**Supplementary Table 1. Demographic and clinical information for human post-mortem tissue cases.** Diagnoses of Parkinson disease were determined clinically using patient histories received from the donors' physicians. Pathological identification of Lewy pathology and dopamine neuron loss in the SNc by brain bank neuropathologists confirmed clinical findings. All Parkinson disease cases were free of other neurological or neuropathological conditions. Age-matched control cases were free of any clinically diagnosed neurological disorders and neuropathological abnormalities.

| Case | Diagnostic | Age | Sex | PMD | Pathological Diagnosis | Cause of Death | Experiment | *SOD1* genotyping |
|---|---|---|---|---|---|---|---|---|
| 1 | PD | 89 | F | 54 | Consistent with Parkinson disease | Parkinson disease | PTM | Performed in this study |
| 2 | PD | 80 | F | 72 | Consistent with Parkinson disease | Parkinson disease | PTM, metallation, pI | Performed in this study |
| 3 | PD | 59 | M | 23 | Consistent with Parkinson disease | Parkinson disease | PTM, metallation, pI | Performed in this study |
| 4 | PD | 84 | M | 34 | Consistent with Parkinson disease | Parkinson disease | PTM, metallation, pI | Performed in this study |
| 5 | PD | 81 | F | 38 | Consistent with Parkinson disease | Parkinson disease | PTM, metallation, pI | Performed in this study |
| 6 | PD | 73 | F | 82 | Consistent with Parkinson disease | Parkinson disease | PTM, metallation, pI | Performed in this study |
| 7 | PD | 82 | M | 28 | Consistent with Parkinson disease | Parkinson disease | PTM, metallation, pI | Performed in this study |
| 8 | PD | 81 | F | 28 | Consistent with Parkinson disease | Parkinson disease | PTM, metallation, pI | Performed in this study |
| 9 | PD | 84 | F | 25 | Consistent with Parkinson disease | Parkinson disease | PTM, metallation, pI | Performed in this study |
| 10 | PD | 80 | M | 29 | Consistent with Parkinson disease | Parkinson disease | PTM, metallation, pI | Performed in this study |
| 11 | PD | 69 | M | 5 | Consistent with Parkinson disease | bronchopneumonia | Metallation, pI | Trist et al., 2018[1] |
| 12 | PD | 74 | M | 16 | Consistent with Parkinson disease | cerebrovascular accident | Metallation, pI | Trist et al., 2018[1] |
| 13 | PD | 75 | M | 9 | Consistent with Parkinson disease | cardiorespiratory failure | Metallation, pI | Trist et al., 2018[1] |
| 14 | PD | 82 | M | 19 | Consistent with Parkinson disease | cardiorespiratory failure | Metallation, pI | Trist et al., 2018[1] |
| 15 | PD | 82 | M | 14 | Consistent with Parkinson disease | cardiorespiratory failure | Metallation, pI | Trist et al., 2018[1] |
| 16 | PD | 82 | F | 3 | Consistent with Parkinson disease | cerebrovascular Alzheimer's disease | Metallation, pI | Trist et al., 2018[1] |
| 17 | PD | 83 | F | 32 | Consistent with Parkinson disease | cardiorespiratory failure | Metallation, pI | Trist et al., 2018[1] |
| 18 | PD | 83 | F | 7 | Consistent with Parkinson disease | pneumonia | Metallation, pI | Trist et al., 2018[1] |
| 19 | PD | 90 | M | 5 | Consistent with Parkinson disease | respiratory failure | Metallation, pI | Trist et al., 2018[1] |
| 20 | Ct | 63 | M | 23 | Control case | colon cancer | PTM, metallation, pI | Performed in this study |
| 21 | Ct | 74 | F | 66 | Braak stage 2 consistent with ageing, control | hospital acquired pneumonia, COPD | PTM, metallation, pI | Performed in this study |
| 22 | Ct | 80 | F | 22 | Consistent with ageing (Tau Braak stage 2) control | stage 4 Lung cancer | PTM, metallation, pI | Performed in this study |
| 23 | Ct | 82 | M | 24 | Ageing process, consistent with Braak stage 2 | | PTM, metallation, pI | Performed in this study |
| 24 | Ct | 82 | M | 47 | control case (very early Alzheimer disease pathology - BNE stage I) with focal amyloid angiopathy | congestive cardiac failure; aortic stenosis | PTM, metallation, pI | Performed in this study |

| | | | | | | | | |
|---|---|---|---|---|---|---|---|---|
| 25 | Ct | 84 | F | 34 | mild pathological ageing changes consistent with HP-tau stage I-II in limbic system; possible control | sepsis; aspiration pneumonia; metastatic breast cancer | PTM, metallation, pI | Performed in this study |
| 26 | Ct | 85 | F | 45 | Ageing process with tau deposition reaching modified Braak (BNE) stage 2 and extensive beta-amyloid plaques. No amyloid angiopathy. Possible control | cancer breast | PTM, metallation, pI | Performed in this study |
| 27 | Ct | 90 | M | 45 | Mild age-related changes (control brain) - AD modified Braak stage + with mild focal amyloid angiopathy | myocardial infarction | PTM, metallation, pI | Performed in this study |
| 28 | Ct | 90 | F | 50 | control brain- mild alzheimer-type changes (modified Braak stage II) and mild amyloid angiopathy | | PTM, metallation, pI | Performed in this study |
| 29 | Ct | 80 | F | 3 | minimal ageing changes, consistent with HP-Tau stage I; childhood poliomyelitis | cancer | PTM, metallation, pI | Performed in this study |
| 30 | Ct | 41 | M | 48 | Control case | cardiorespiratory failure | Metallation, pI | Trist et al., 2018[1] |
| 31 | Ct | 49 | M | 47 | Control case | cardiorespiratory failure | Metallation, pI | Trist et al., 2018[1] |
| 32 | Ct | 65 | M | 14.5 | Control case | cardiorespiratory failure | Metallation, pI | Trist et al., 2018[1] |
| 33 | Ct | 74 | M | 16.5 | Control case | toxicity | Metallation, pI | Trist et al., 2018[1] |
| 34 | Ct | 84 | F | 6 | Control case | cardiorespiratory failure | Metallation, pI | Trist et al., 2018[1] |
| 35 | Ct | 87 | F | 24 | Control case | acute peritonitis | Metallation, pI | Trist et al., 2018[1] |
| 36 | Ct | 89 | M | 22 | Control case | cardiorespiratory failure | Metallation, pI | Trist et al., 2018[1] |
| 37 | Ct | 89 | F | 23 | Control case | metastatic adenocarcinoma | Metallation, pI | Trist et al., 2018[1] |
| 38 | Ct | 92 | F | 5 | Control case | pancytopaenia | Metallation, pI | Trist et al., 2018[1] |
| 39 | Ct | 102 | F | 5 | Control case | acute renal failure | Metallation, pI | Trist et al., 2018[1] |
| 40 | Ct | 103 | M | 20 | Control case | acute myocardial infarct | Metallation, pI | Trist et al., 2018[1] |
| 41 | Ct | 94 | F | 6 | Control case | cardiorespiratory failure | Metallation, pI | Trist et al., 2018[1] |
| 42 | Ct | 54 | M | 10 | Control case | cardiorespiratory failure | Metallation, pI | Trist et al., 2018[1] |

**Abbreviations**: AD, Alzheimer's disease; COPD, chronic obstructive pulmonary disorder; F, female; M, male; PD, Parkinson disease; PMD, post-mortem delay.

**Supplementary Table 2. Demographic statistics for diagnostic groups.**

| Variable | PD | Control | Significantly different? |
|---|---|---|---|
| *n* | 19 | 23 | N/A |
| Sex (M:F) | 10:9 | 10:13 | N.S. ($p$ = 0.76)[a] |
| Age (years) | 79.6 ± 7.1 | 79.7 ± 15.9 | N.S. ($p$ = 0.98)[b] |
| [range] | 59 - 90 | 41 - 103 | |
| PMI (hrs) | 27.5 ± 21.9 | 26.4 ± 17.9 | N.S. ($p$ = 0.85)[b] |

**Footnotes:** [a] Chi-square test, [b] Unpaired *t*-test

**Abbreviations:** F, female; M, male; N/A, not applicable; N.S., not significant; PD, Parkinson disease; PMI, post-mortem interval.

**Supplementary Table 3. Genetic screening of the SOD1 gene in PD patients and controls.** DNA was extracted from human brain tissue using the Qiagen DNeasy DNA extraction kit, following standard protocol. All five exons of SOD1, and at least 10bp of flanking sequence were sequenced using PCR amplification and Sanger sequencing. All results were blindly and independently analysed by two team members. A non-coding 'A' deletion (intron 5, c.360+90DelA, rs3216079) was found in two PD cases. This deletion is present in multiple control cohorts (e.g. Turkish Genome Project, Gene ID ENSG00000142168), and not of clinical significance. Remaining PD and control cases were genotyped previously in Trist et al., 2018[1].

| Case # | Diagnostic | DNA conc. (ug/uL) | A260/ A280 | SOD1 exon 1 | SOD1 exon 2 | SOD1 exon 3 | SOD1 exon 4 | SOD1 exon 5 |
|---|---|---|---|---|---|---|---|---|
| 1 | PD | 53.3 | 1.89 | negative | negative | negative | negative | negative |
| 2 | PD | 54.7 | 1.91 | negative | negative | negative | negative | negative |
| 3 | PD | 53.4 | 1.98 | negative | negative | negative | negative | negative |
| 4 | PD | 72.5 | 2.02 | negative | negative | negative | negative | negative |
| 5 | PD | 39 | 1.89 | negative | negative | negative | rs3216079 | negative |
| 6 | PD | 89.2 | 1.98 | negative | negative | negative | negative | negative |
| 7 | PD | 102.4 | 1.86 | negative | negative | negative | negative | negative |
| 8 | PD | 63.2 | 2 | negative | negative | negative | rs3216079 | negative |
| 9 | PD | 47.3 | 1.98 | negative | negative | negative | negative | negative |
| 10 | PD | 81.7 | 1.93 | negative | negative | negative | negative | negative |
| 20 | Control | 37.9 | 1.92 | negative | negative | negative | negative | negative |
| 21 | Control | 59.2 | 2.06 | negative | negative | negative | negative | negative |
| 22 | Control | 70.7 | 2.02 | negative | negative | negative | negative | negative |
| 23 | Control | 78 | 1.98 | negative | negative | negative | negative | negative |
| 24 | Control | 49.6 | 1.89 | negative | negative | negative | negative | negative |
| 25 | Control | 81.9 | 1.96 | negative | negative | negative | negative | negative |
| 26 | Control | 40 | 1.93 | negative | negative | negative | negative | negative |
| 27 | Control | 44.9 | 1.89 | negative | negative | negative | negative | negative |
| 28 | Control | 73 | 1.98 | negative | negative | negative | negative | negative |
| 29 | Control | 86.7 | 1.9 | negative | negative | negative | negative | negative |

**Abbreviations:** A260, absorbance 260nm; A280, absorbance 280nm, SOD1, superoxide dismutase 1.

**Supplementary Table 4. SOD1 PTMs in post-mortem PD and control substantia nigra.** SOD1 protein was isolated from post-mortem tissues using our published immunoprecipitation method [2], which we have demonstrated does not significantly alter SOD1 PTMs. Peptides were also prepared as previously described [2] and bottom-up proteomic mass spectrometry performed on a Q Exactive HF-X Hybrid Quadrupole-Orbitrap Mass Spectrometer operated in data-independent acquisition mode. Data were analyzed using Spectronaut proteomics software (Version 18, Biognosys, Schlieren, Zurich, Switzerland), with the threshold for statistical significance adjusted using the Bonferroni correction method to counteract the higher type 1 error rate resulting from using multiple *t*-tests during data analysis ($p < 0.05$/# pairings, 6 pairings, new threshold; $p < 0.00833$)[3]. Full methods details are presented in the main methods section. Amino acid residues are identified using one letter code. Sample size = 10/group. Atypical PTMs (top) are separated from physiological PTMs (bottom).

| Modification | Modified residues (all groups) | Altered PD vs Ct SN? | Test statistics (p, Q, # ratios) | Alteration present in OCx? |
|---|---|---|---|---|
| Oxidation (H) | H43, H46, H48, H63, H80, H110, H120 | H80 (6.8-fold increase) | 0.00017, 0.00073, 68 | No |
| Oxidation (W) | W32 | No | N/A | N/A |
| Kynurenine (W) | W32 | W32 (1.5-fold increase) | 0.0026, 0.0035, 80 | No |
| Hydroxykynurenine (W) | W32 | No | N/A | N/A |
| Dioxidation (W) | W32 | No | N/A | N/A |
| Nitration (W) | W32 | No | N/A | N/A |
| Carboxymethyllysine (K) | K9, K36, K91 | K36 (3.7-fold increase) | 0.00022, 0.0037, 34 | No |
| Glycation (R, K) | K9, K23, R69, K91, R115, K122, K128, K136 | K9 (2.6-fold increase) | 0.00006, 0.00013, 48 | No |
| | | K23 (2.6-fold increase) | 0.00006, 0.00013, 48 | No |
| | | K91 (1.9-fold increase) | 0.000018, 0.000058, 64 | No |
| | | R115 (1.9-fold increase) | 0.000018, 0.000058, 64 | No |
| | | K122 (2.1-fold increase) | 0.0013, 0.0018, 30 | No |
| Gloxal AGE (R) | R69, R115 | No | N/A | N/A |
| Acetylation (K) | K3, K9, K23, K91, K122 | No | N/A | N/A |
| Succinylation (K) | K3, K9, K23, K30, K36 | No | N/A | N/A |
| Phosphorylation (S, T) | T2, S25, T39, T54, T58, S59, S68, T88, S102, T116, S142 | No | N/A | N/A |

| | | | | |
|---|---|---|---|---|
| Deamidation (N, Q) | Q15, N19, Q22, N26, N53, N65, N86, N131, N139, Q153 | No | N/A | |
| Ubiquitylation (GlyGly footprint; K) | K9, K91 | No | N/A | N/A |
| Glycosylation (N, S, T) | T2, N19, N53, T54, T58, S59, T88, S98, S102, S105, S107, T116, N131 | S98 (79.1-fold decrease) | 0.0019, 0.0037, 12 | No |
| | | N131 (5.1-fold decrease) | 0.0014, 0.0034, 62 | No |
| Acetylglucosamine (N, S, T) | N19, S25, N26, S98, S102, S105, S107, T116, N131, T137 | S25 (1.6-fold decrease) | 0.0011, 0.0041, 56 | No |
| | | N26 (1.6-fold decrease) | 0.0011, 0.0041, 56 | No |
| | | N131 (4.2-fold decrease) | 0.00038, 0.0018, 64 | No |

**Abbreviations:** AGE, advanced-glycation end-product; Ct, control; N/A, not applicable; OCx, occipital cortex; PD, Parkinson disease; SN, substantia nigra.

**Supplementary Table 5. SOD1 PTMs in post-mortem PD and control occipital cortex.** Protein preparation, as well as mass spectrometry data acquisition and analysis, were performed as described in **Supplementary Table 4.** Amino acid residues are identified using three letter code. Sample size = 10/group.

| **Modification** | **Modified residues (all groups)** | **Fold change in PD OCx** | **Test statistics (*p*, Q, # ratios)** |
|---|---|---|---|
| Deamidation (N, Q) | Q15, N19, Q22, N26, N53, N65, N86, N131, N139, Q153 | Q15 (1.8-fold increase) | 0.00038, 0.00099, 80 |
| | | N19 (2.9-fold increase) | 0.000058, 0.00023, 72 |
| | | N26 (1.6-fold increase) | 0.00062, 0.0015, 80 |
| | | N86 (1.9-fold increase) | 0.000085, 0.00033, 78 |
| | | N131 (2.5-fold increase) | 0.0000052, 0.000034, 80 |
| Glycosylation (N, S, T) | T2, N19, N53, T54, T58, S59, N86, T88, S98, S102, S105, S107, T116, N131, S134, T135, T137 | T54 (3.9-fold increase) | 0.00034, 0.00063, 32 |
| | | T86 (1.9-fold increase) | 0.0015, 0.002, 62 |
| | | T88 (2-fold increase) | 0.0005, 0.00076, 62 |
| | | S98 (2-fold increase) | 0.0005, 0.00076, 62 |
| | | S102 (2-fold increase) | 0.0005, 0.00076, 62 |
| | | S105 (2-fold increase) | 0.0005, 0.00076, 62 |
| | | S107 (6.7-fold increase) | 0.00025, 0.00054, 34 |
| | | T116 (2.4-fold increase) | 0.00056, 0.00082, 20 |
| | | N131 (4.2-fold increase) | 0.0017, 0.0022, 72 |
| | | S134 (3.5-fold increase) | 0.000019, 0.00008, 28 |
| | | T135 (3.5-fold increase) | 0.000019, 0.00008, 28 |
| | | T137 (3.5-fold increase) | 0.000019, 0.00008, 28 |
| Acetylglucosamine (N, S, T) | N19, S25, N26, S98, S102, S105, S107, T116, N131, T137 | S98 (1.6-fold increase) | 0.00031, 0.0011, 62 |

**Abbreviations:** Ct, control; N/A, not applicable; OCx, occipital cortex; PD, Parkinson disease.

**Supplementary Table 6. Altered SOD1 PTMs in the SOCK mouse midbrain compared with healthy aged human SNc.** Mice were 6 months-of-age. Protein preparation, as well as mass spectrometry data acquisition and analysis, were performed as described in **Supplementary Table 4.** Amino acid residues are identified using three letter code. Sample size = 10/group. Atypical PTMs (top) are separated from physiological PTMs (bottom).

| Modification | Modified residues (all groups) | Altered SOCK mouse vs aged human SN ($Log_2$(ratio)) | Test statistics (p, Q, # ratios) | Alteration also present in h*SOD1^WT^* mice? |
|---|---|---|---|---|
| Oxidation (H) | H43, H46, H48, H63, H71, H80, H110, H120 | H46 (0.706) | 0.0020, 0.00068, 76 | No |
| | | H48 (0.803) | 0.016, 0.0045, 58 | No |
| | | H71 (3.469) | 0.0033, 0.0010, 52 | No |
| | | H80 (-0.782) | 0.00012, 5.22E-05, 66 | Yes |
| | | H110 (2.347) | 1.72E-07, 9.83E-08, 80 | Yes |
| | | H120 (-1.208) | 7.53E-06, 3.76E-06, 78 | Yes |
| Oxidation (W) | W32 | No | N/A | N/A |
| Kynurenine (W) | W32 | No | N/A | N/A |
| Hydroxykynurenine (W) | W32 | No | N/A | N/A |
| Dioxidation (W) | W32 | No | N/A | N/A |
| Nitration (W) | - | N/A | N/A | N/A |
| Carboxymethyllysine (K) | K9, K3, K91, K128, K122, K23 | K91 (-1.445) | 5.41E-06, 5.41E-06, 80 | Yes |
| | | K128 (2.722) | 3.40E-19, 1.36E-18, 74 | Yes |
| | | K122 (2.947) | 2.30E-20, 1.84E-19, 72 | Yes |
| | | K9 (-1.024) | 1.82E-07, 2.92E-07, 76 | Yes |
| | | K3 (-1.053) | 2.34E-07, 3.12E-07, 76 | Yes |
| | | K23 (2.172) | 0.002438553, 0.001625702, 46 | No |
| Glycation (R, K) | K3, K23, K30, K36, R69, K75, R79, K91, K128, K136 | K3 (-0.790) | 1.39E-08, 2.23E-08, 76 | Yes |
| | | K23 (-1.399) | 0.00018765, 0.00015012, 80 | Yes |
| | | K91 (2.106) | 5.76E-12, 1.53E-11, 56 | Yes |
| | | K128 (2.033) | 5.16E-12, 1.53E-11, 62 | Yes |
| | | K136 (2.033) | 2.66E-13, 2.12E-12, 64 | Yes |
| Gloxal AGE (R) | R69, R115, R143 | No | N/A | N/A |
| Acetylation (K) | K9, K23, K30, K36, K128 | K9 (-1.811) | 4.72E-08, 1.57E-08, 28 | No |
| | | K23 (-1.811) | 4.72E-08, 1.57E-08, 28 | No |

| | | | | |
|---|---|---|---|---|
| | | K30 (-2.112) | 3.41E-12, 2.27E-12, 54 | Yes |
| | | K36 (1.566) | 1.60E-17, 3.19E-17, 78 | Yes |
| | | K128 (0.957) | 3.86E-09, 1.93E-09, 80 | Yes |
| Succinylation (K) | K91, K30, K23, K9, K75, K70, K3 | K91 (-0.763) | 0.000294989, 0.000294989, 80 | No |
| | | K30 (-1.144) | 2.87E-15, 2.87E-15, 76 | Yes |
| | | K23 (6.200) | 4.52E-23, 4.52E-23, 62 | Yes |
| | | K9 (6.182) | 1.09E-22, 1.09E-22, 62 | Yes |
| | | K75 (4.862) | 2.66E-28, 2.66E-28, 52 | Yes |
| | | K70 (4.862) | 2.66E-28, 2.66E-28, 52 | Yes |
| | | K3 (6.112) | 1.31E-22, 1.31E-22, 60 | Yes |
| Phosphorylation (S, T) | S34, S25, T54, S134, S68, S59 | S34 (1.424) | 5.26E-14, 5.26E-14, 60 | Yes |
| | | S25 (1.424) | 5.26E-14, 5.26E-14, 60 | Yes |
| | | T54 (-0.898) | 0.003190631, 0.003190631, 62 | Yes |
| | | S134 (0.871) | 0.000328369, 0.000328369, 60 | Yes |
| | | S68 (0.669) | 7.82E-09, 7.82E-09, 50 | Yes |
| | | S59 (0.669) | 7.82E-09, 7.82E-09, 50 | Yes |
| Deamidation (N, Q) | Q22, N26, Q15, N19, N53, N139, N131 | Q22 (-0.764) | 7.38E-05, 6.15E-05, 54 | No |
| | | N26 (-1.329) | 2.98E-16, 7.46E-16, 80 | Yes |
| | | Q15 (0.857) | 8.38E-07, 1.20E-06, 80 | Yes |
| | | N19 (1.863) | 1.57E-23, 1.57E-22, 80 | Yes |
| | | N53 (0.754) | 4.20E-13, 8.41E-13, 78 | Yes |
| | | N139 (3.288) | 3.92E-05, 3.92E-05, 64 | Yes |
| | | N131 (1.185) | 9.12E-22, 3.04E-21, 78 | Yes |
| Ubiquitylation (GlyGly footprint; K) | K9, K23, K136, K91, K128, K3 | K9 (-1.565) | 1.01E-07, 2.90E-08, 78 | No |
| | | K23 (1.128) | 2.36E-13, 1.18E-13, 64 | Yes |
| | | K136 (-2.017) | 1.64E-06, 4.09E-07, 34 | No |
| | | K91 (-0.770) | 0.000489975, 9.80E-05, 30 | No |
| | | K128 (-0.688) | 0.057710929, 0.008878605, 64 | No |
| | | K3 (1.834) | 2.33E-20, 1.55E-20, 60 | Yes |
| Glycosylation (N, S, T) | T2, N19, S25, N26, T39, T58, N86, T88, | S25 (-1.513) | 0.000379713, 0.000350504, 34 | No |
| | | N26 (-1.513) | 0.000379713, 0.000350504, 34 | No |

| | | | | |
|---|---|---|---|---|
| | S98, S102, S105, S107, T116, N131 | T39 (1.820) | 2.73E-13, 3.27E-12, 38 | Yes |
| | | T58 (-1.262) | 0.000596319, 0.00051113, 64 | Yes |
| | | N86 (-1.679) | 5.92E-06, 1.02E-05, 68 | No |
| | | T88 (-1.679) | 5.92E-06, 1.02E-05, 68 | No |
| | | S98 (-1.246) | 1.21E-05, 1.61E-05, 80 | Yes |
| | | S102 (-1.246) | 1.21E-05, 1.61E-05, 80 | Yes |
| | | S105 (-1.179) | 2.44E-05, 2.66E-05, 80 | Yes |
| | | S107 (-4.093) | 4.92E-12, 2.95E-11, 78 | Yes |
| | | T116 (-0.989) | 1.50E-05, 1.80E-05, 74 | Yes |
| | | N131 (0.883) | 3.97E-10, 1.59E-09, 76 | Yes |
| Acetylglucosamine (N, S, T) | T88, N86, S105, S102, S98, T39, S131, T58 | T88 (-1.182) | 6.48E-05, 4.86E-05, 74 | No |
| | | N86 (-1.182) | 6.48E-05, 4.86E-05, 74 | No |
| | | S105 (-2.534) | 1.88E-05, 2.82E-05, 46 | No |
| | | S102 (0.862) | 3.02E-05, 3.02E-05, 64 | Yes |
| | | T39 (1.222) | 0.001589265, 0.00105951, 30 | Yes |
| | | S98 (-0.603) | 0.001890488, 0.001134293, 80 | No |
| | | N131 (1.364) | 1.56E-16, 9.35E-16, 80 | Yes |
| | | T57 (-1.713) | 1.94E-10, 5.81E-10, 64 | Yes |

**Abbreviations:** AGE, advanced glycation end-product; N/A, not applicable; SN, substantia nigra.

**Supplementary Table 7. Altered SOD1 PTMs in the h*SOD1*$^{WT}$ mouse midbrain compared with healthy aged human SNc.** Mice were 6 months of age. Protein preparation, as well as mass spectrometry data acquisition and analysis, were performed as described in **Supplementary Table 4.** Amino acid residues are identified using three letter code. Sample size = 10/group. Atypical PTMs (top) are separated from physiological PTMs (bottom).

| **Modification** | **Modified residues (all groups)** | **Altered hSOD1WT mouse vs aged human SN (Log2(ratio))** | **Test statistics (p, Q, # ratios)** |
|---|---|---|---|
| Oxidation (H) | H43, H46, H48, H63, H71, H80, H110, H120 | H43 (-1.012) | 0.0054, 0.0015, 84 |
| | | H46 (-1.014) | 0.0054, 0.0015, 84 |
| | | H43 (-1.012) | 0.0054, 0.0015, 84 |
| | | H63 (-5.241) | 6.29E-16, 1.26E-15, 36 |
| | | H80 (-0.792) | 0.00013, 6.25E-05, 76 |
| | | H110 (1.977) | 9.26E-14, 1.23E-13, 84 |
| | | H120 (-4.051) | 4.90E-17, 1.96E-16, 76 |
| Oxidation (W) | W32 | No | N/A |
| Kynurenine (W) | W32 | No | N/A |
| Hydroxykynurenine (W) | W32 | No | N/A |
| Dioxidation (W) | W32 | No | N/A |
| Nitration (W) | - | N/A | N/A |
| Carboxymethyllysine (K) | K9, K3, K91, K128, K122, K23 | K91 (-1.835) | 5.29E-08, 5.29E-08, 84 |
| | | K128 (2.744) | 7.48E-19, 9.97E-19, 78 |
| | | K122 (2.969) | 3.95E-20, 7.91E-20, 76 |
| | | K9 (-0.72) | 1.40E-05, 1.12E-05, 84 |
| | | K3 (-0.744) | 1.73E-05, 1.15E-05, 84 |
| Glycation (R, K) | K3, K23, K30, K36, R69, K75, R79, K91, K128, K136 | K9 (-1.216) | 0.000412254, 0.000471147, 72 |
| | | K91 (2.585) | 1.01E-10, 2.02E-10, 60 |
| | | K128 (1.914) | 9.31E-16, 2.48E-15, 74 |
| | | K136 (2.043) | 4.60E-16, 1.84E-15, 70 |
| Gloxal AGE (R) | R69, R115, R143 | N/A | N/A |
| Acetylation (K) | K9, K23, K30, K36, K128 | K30 (-1.181) | 3.53E-05, 2.35E-05, 50 |
| | | K36 (1.462) | 5.14E-11, 1.03E-10, 84 |
| | | K128 (0.856) | 5.33E-06, 4.27E-06, 84 |
| Succinylation (K) | K91, K30, K23, K9, K75, K70, K3 | K30 (-0.911) | 3.98E-11, 2.27E-11, 80 |
| | | K23 (6.051) | 8.94E-30, 3.57E-29, 64 |
| | | K9 (6.023) | 1.80E-27, 2.11E-27, 62 |
| | | K75 (5.484) | 2.11E-27, 2.11E-27, 54 |
| | | K70 (5.484) | 2.11E-27, 2.11E-27, 54 |
| | | K3 (5.914) | 4.33E-27, 3.47E-27, 60 |

| | | | |
|---|---|---|---|
| Phosphorylation (S, T) | S34, S25, T54, S134, S68, S59 | S34 (1.828) | 3.61E-14, 2.41E-14, 72 |
| | | S25 (1.828) | 3.61E-14, 2.41E-14, 72 |
| | | T54 (-1.161) | 0.00056158, 0.000102106, 64 |
| | | S134 (1.732) | 7.06E-07, 2.02E-07, 62 |
| | | S102 (-1.084) | 3.81E-05, 8.47E-06, 56 |
| | | S68 (0.878) | 9.96E-12, 3.98E-12, 58 |
| | | S59 (0.878) | 9.96E-12, 3.98E-12, 58 |
| | | T39 (0.722) | 8.30E-08, 2.77E-08, 70 |
| Deamidation (N, Q) | Q22, N26, Q15, N19, N53, N139, N131 | Q153 (1.465) | 2.28E-05, 9.75E-06, 84 |
| | | N26 (0.820) | 8.18E-13, 8.18E-13, 84 |
| | | Q15 (1.102) | 1.78E-12, 1.53E-12, 84 |
| | | N19 (1.012) | 5.70E-11, 4.28E-11, 84 |
| | | N53 (1.079) | 6.35E-20, 1.27E-19, 84 |
| | | N139 (0.989) | 1.86E-18, 2.23E-18, 82 |
| | | N131 (0.989) | 1.86E-18, 2.23E-18, 82 |
| Ubiquitylation (GlyGly footprint; K) | K9, K23, K136, K91, K128, K3 | K23 (1.498) | 1.05E-17, 1.05E-17, 70 |
| | | K9 (0.592) | 0.012916516, 0.00774991, 84 |
| | | K3 (2.781) | 9.28E-22, 1.39E-21, 72 |
| Glycosylation (N, S, T) | T2, N19, S25, N26, T39, T58, N86, T88, S98, S102, S105, S107, T116, N131 | T39 (2.080) | 8.24E-14, 4.94E-13, 36 |
| | | T58 (-1.361) | 0.000310903, 0.000532976, 66 |
| | | S98 (-0.857) | 0.001861984, 0.002234381, 84 |
| | | S102 (-0.857) | 0.001861984, 0.002234381, 84 |
| | | S105 (-0.793) | 0.003451079, 0.003764813, 84 |
| | | S107 (-2.045) | 2.69E-05, 8.07E-05, 82 |
| | | T116 (-0.738) | 0.000727347, 0.001091021, 78 |
| | | N131 (0.813) | 1.21E-07, 4.83E-07, 80 |
| Acetylglucosamine (N, S, T) | T88, N86, S105, S102, S98, T39, S131, T58 | S102 (1.165) | 1.96E-08, 2.36E-08, 74 |
| | | S98 (1.165) | 1.96E-08, 2.36E-08, 74 |
| | | T39 (1.207) | 0.00158075, 0.000862227, 36 |
| | | N131 (1.649) | 4.76E-21, 1.43E-20, 84 |
| | | T58 (-1.650) | 2.16E-09, 4.33E-09, 70 |

**Supplementary Table 8. Primary and secondary antibody details for all methods used in this study.**

| Antibody | Source and catalogue # | Type | Host | Species reactivity | Immunogen | Application | Dilution |
|---|---|---|---|---|---|---|---|
| SOD1 UβB | StressMarq Biosciences, British Columbia, Canada (SPC-205) | P | Rb | Ms, Hu, Rat | N-terminal region, SOD1 protein with unfolded β-barrel | IF | 1:300 |
| SOD1 EDI | StressMarq Biosciences, British Columbia, Canada (SPC-206) | P | Rb | Ms, Hu, Rat | C-terminal region of human SOD1, exposed dimer interface | IF | 1:500 |
| Pan-SOD1 | Enzo Biochem, New York, USA (ADI-SOD-100) | P | Rb | Ms, Hu, Rat | Native human Cu/Zn SOD | IF | 1:1000 |
| | | | | | | WB | 1:2000 |
| | | | | | | IP | 12.5ug/mg beads |
| TH | Merck Millipore, California, USA (AB152) | P | Rb | Ms, Hu, Rat | Denatured tyrosine hydroxylase from rat pheochromocytoma (denatured by SDS). | IHC | 1:5000 |
| CCS | Jonsson et al., 2006 [4] | P | Rb | Ms, Hu | Amino acids 252–270 of human CCS | WB | 1:2000 |
| GAPDH | Sigma Aldrich, St Louis, Missouri, USA (G9545) | P | Rb | Ms, Hu, Rat | Synthetic peptide corresponding to amino acids of mouse GAPDH, conjugated to KLH via an N-terminal cysteine residue | WB | 1:1000 |
| TH | Abcam, Cambridge, United Kingdom (ab76442) | P | Ch | Ms | Synthetic peptides corresponding to sequences shared between murine (P24529) and human (P07101) TH | IF | 1:300 |
| NeuN | Merck Millipore, California, USA (ABN91) | P | Ch | Rb, Ms | First 97 amino acids from the N-terminal region of murine NeuN | IF | 1:800 |
| GFAP | Novus Biologicals, Colorado, USA (NBP1-05198) | P | Ch | Ms, Hu, Rat | Recombinant full length human GFAP isotype 1 | IF | 1:5000 |
| Iba1 | Invitrogen, Massachusetts, USA (MA5-38266) | M | Rat | Ms, Hu, Rat | Iba1 C-terminus | IF | 1:2000 |
| pS129 α-synuclein | Abcam, Cambridge, United Kingdom (ab51253) | M | Rb | Ms, Hu, Rat | N/A | IF | 1:2000 |

| | | | | | | | |
|---|---|---|---|---|---|---|---|
| ISL1 | Abcam, Cambridge, United Kingdom (ab109517) | M | Rb | Ms, Hu, Rat | N/A | IF | 1:1000 |
| ChAT | Sigma Aldrich, St Louis, Missouri, USA (AB144P) | P | Gt | Ms, Hu, Rat | Human placental ChAT | IF | 1:500 |
| Biotinylated goat anti-rabbit IgG (H+L) | Vector Laboratories, California, USA (BA-1000) | P | Gt | Rb | Rabbit IgG (H+L) | IHC | 1:200 |
| Fab fragment IgG (Alexa Fluor® 488) | Jackson ImmunoResearch Laboratories, Pennsylvania, USA (103-547-008) | P | Gt | Ch | IgY (IgG), Fc fragment specific | IF | 1:400 |
| Fab fragment IgG (Alexa Fluor® 647) | Jackson ImmunoResearch Laboratories, Pennsylvania, USA (111-607-003) | P | Gt | Rb | IgG (H+L) | IF | 1:400 |

**Abbreviations:** Ch, chicken; EDI, exposed dimer interface; Gt, goat; Hu, human; IF, immunofluorescence; IHC, immunohistochemistry; IP, immunoprecipitation; M, monoclonal; Ms, mouse; NeuN, neuronal nuclear protein; P, polyclonal; Rb, rabbit; Sh, sheep; SOD1, superoxide dismutase 1; TH, Tyrosine hydroxylase; UβB, unfolded beta-barrel.

**Supplementary Table 9. Sample size of mouse groups in each genotype and age cohort.** Group sizes expressed as a ratio of the number of male to female mice. Total number of mice is represented in brackets.

| Age (mo) | Wildtype | *Ctr1*$^{+/-}$ | h*SOD1*$^{WT}$ | SOCK |
|---|---|---|---|---|
| 1.5 | 3:2 (5) | 3:5 (8) | 0:3 (3) | 4:3 (7) |
| 3 | 8:8 (16) | 9:4 (13) | 0:7 (7) | 7:6 (13) |
| 6 | 8:8 (16) | 9:7 (16) | 0:10 (10) | 5:7 (12) |
| 12 | 5:9 (14) | 6:4 (10) | 3:6 (9) | 7:6 (13) |

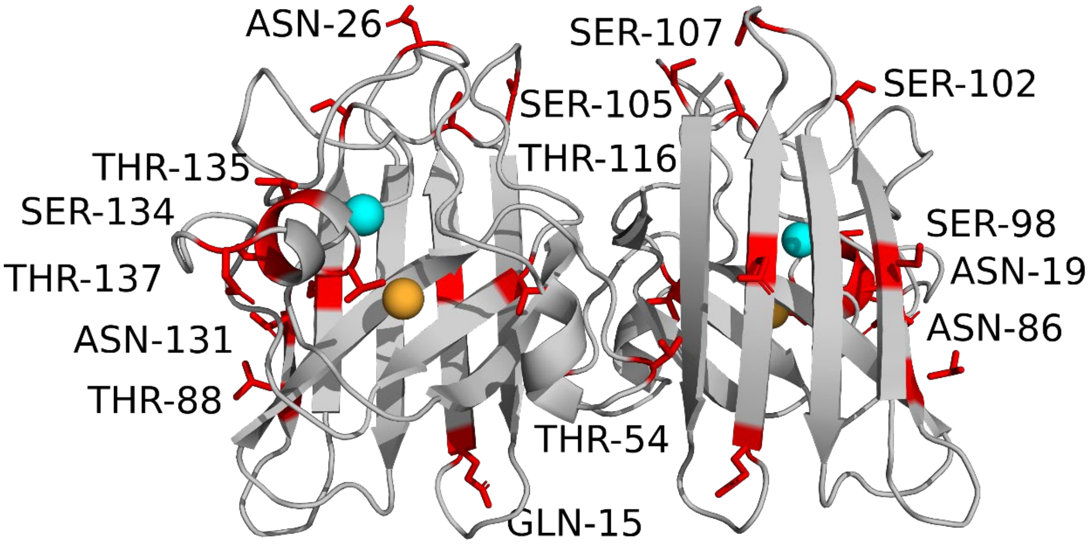


**Supplementary Fig. 1 Altered SOD1 post-translational modifications in the post-mortem PD OCx.** A significant increase in glycosylation of T54, T86, T88, Ser98, Ser102, Ser105, Ser107, T116, Asn131, Ser134, T135 and T137, as well as in deamidation of Gln15, Asn19, Asn26, Asn86 and Asn131, was identified in the OCx of PD patients compared with controls. Residues are labelled using Tee letter amino acid codes with their side chains highlighted in red. Complete details of statistical analyses used to determine PTM alterations are presented in **Supplementary Table 4**.

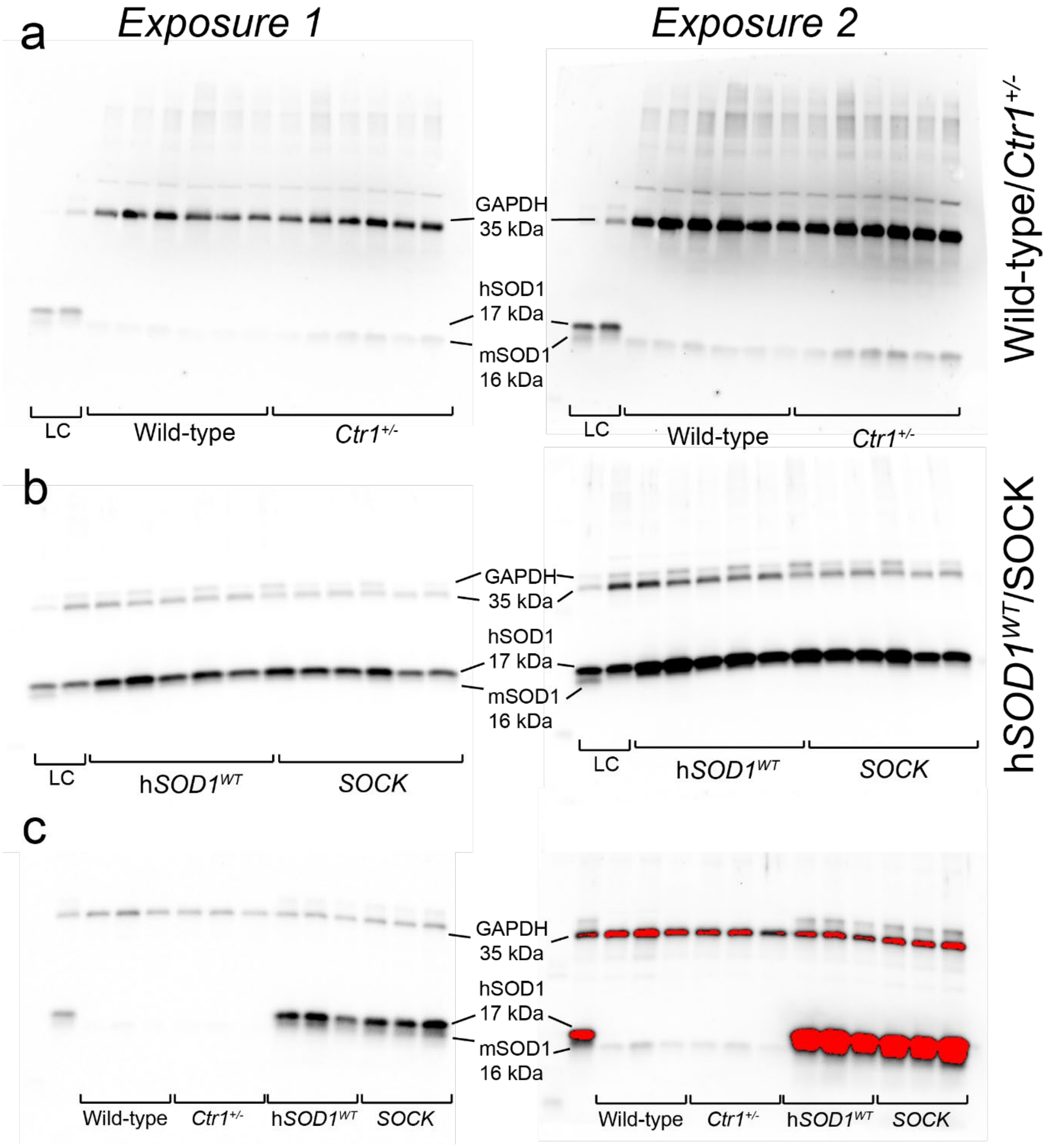


**Supplementary Fig. 2 Representative SOD1 and GAPDH immunoblots from midbrain tissues of all four mouse genotypes.** Extracts from wild-type and *Ctr1*$^{+/-}$ mice (**a**) were run separately to those from h*SOD1*$^{WT}$ and SOCK animals (**b**), given their inclusion in the same gel was technically difficult due to the large differences in SOD1 protein expression between these groups (**c**). A common loading control (LC) made from pooled midbrain, cortex and liver extracts was included on all gels to enable standardization and estimation of protein expression differences between these groups. Blots were probed simultaneously for SOD1 (human isoform; 16 kDa, mouse isoform; 17 kDa) and GAPDH (two bands; 35 and 36 kDa) given their distinct molecular weights (**c**). This was performed in a second round of immunoblotting following initial probing of membranes for CCS (35 kDA), which was successfully stripped prior to SOD1 and GAPDH immunoblotting, as evidenced by the lack of chemiluminescent signal at the molecular weight of CCS (35 kDa) on blots that were stripped, washed, re-probed with HRP-conjugated anti-rabbit secondary antibody and re-imaged with ECL substrate. Full antibody details are listed in **Supplementary Table 8**. Blots were imaged twice at exposures tailored to maximize SOD1 (both isoforms) and GAPDH (both bands) chemiluminescent signals without saturation. Representative blots constitute extracts from midbrain tissues.

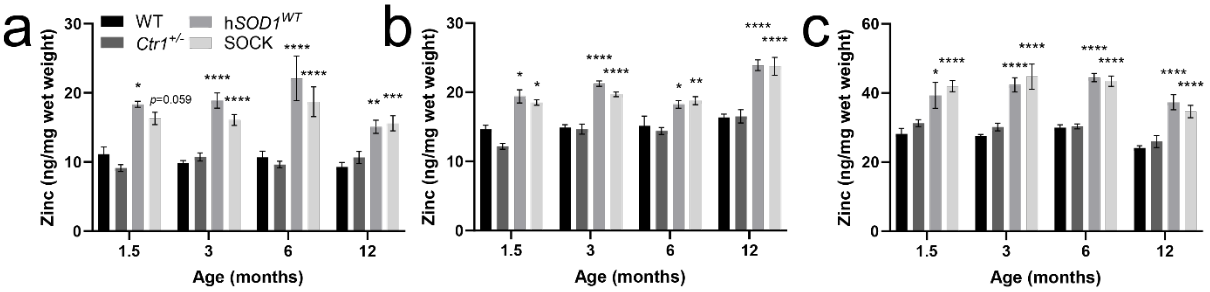


**Supplementary Fig. 3 Zinc levels in the midbrain (a), cortex (b) and liver (c) of all 4 mouse genotypes.** Measurements were obtained using inductively coupled plasma-mass spectrometry (ICP-MS) and differences identified by a two-way ANOVA with Dunnet's multiple comparisons post hoc *t* tests. Sample size per genotype per age: 1.5 month, $n$ = 3-8; 3 month, $n$ = 7-15; 6 month, $n$ = 10-16; 12 month, $n$ = 9-14. Data represents mean ± SEM. *$p < 0.05$, **$p < 0.01$, ***$p < 0.001$, ****$p < 0.0001$. Abbreviations: WT, wild-type.

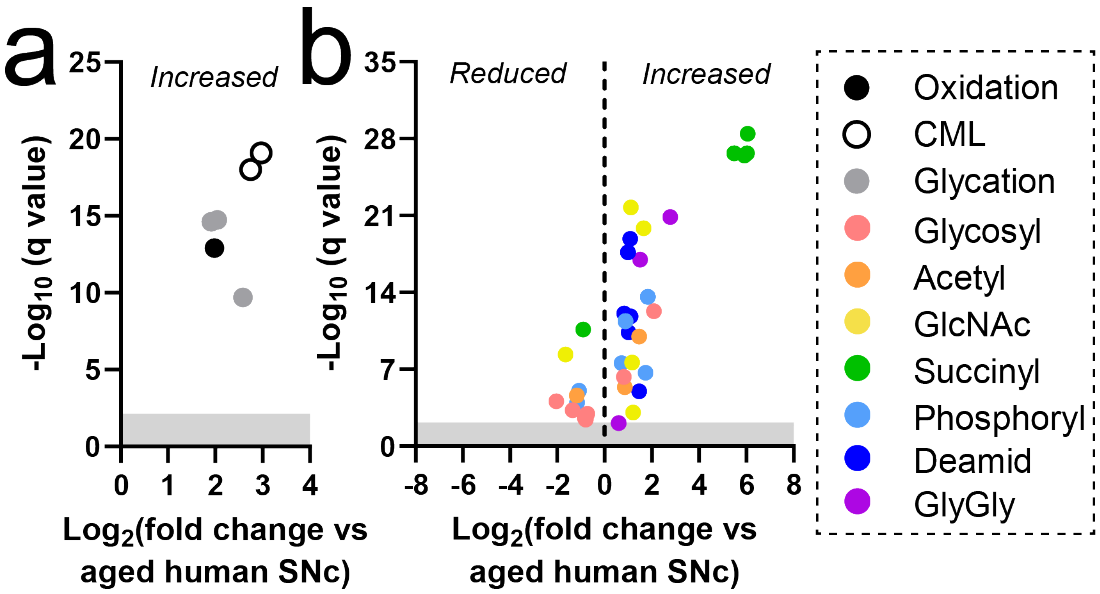


**Supplementary Fig. 4 Altered SOD1 post-translational modifications in the hSOD1$^{WT}$ mouse midbrain.** Complete details of PTM alterations and statistical tests are presented in **Supplementary Table 8**. Abbreviations: CML, carboxymethyllysine; Glycosyl, glycosylation; Acetyl, acetylation; GlcNAc, acetylglucosamine; Succinyl, succinylation; Phosphoryl, phosphorylation; Deamid, deamidation; GlyGly, ubiquitin footprint.

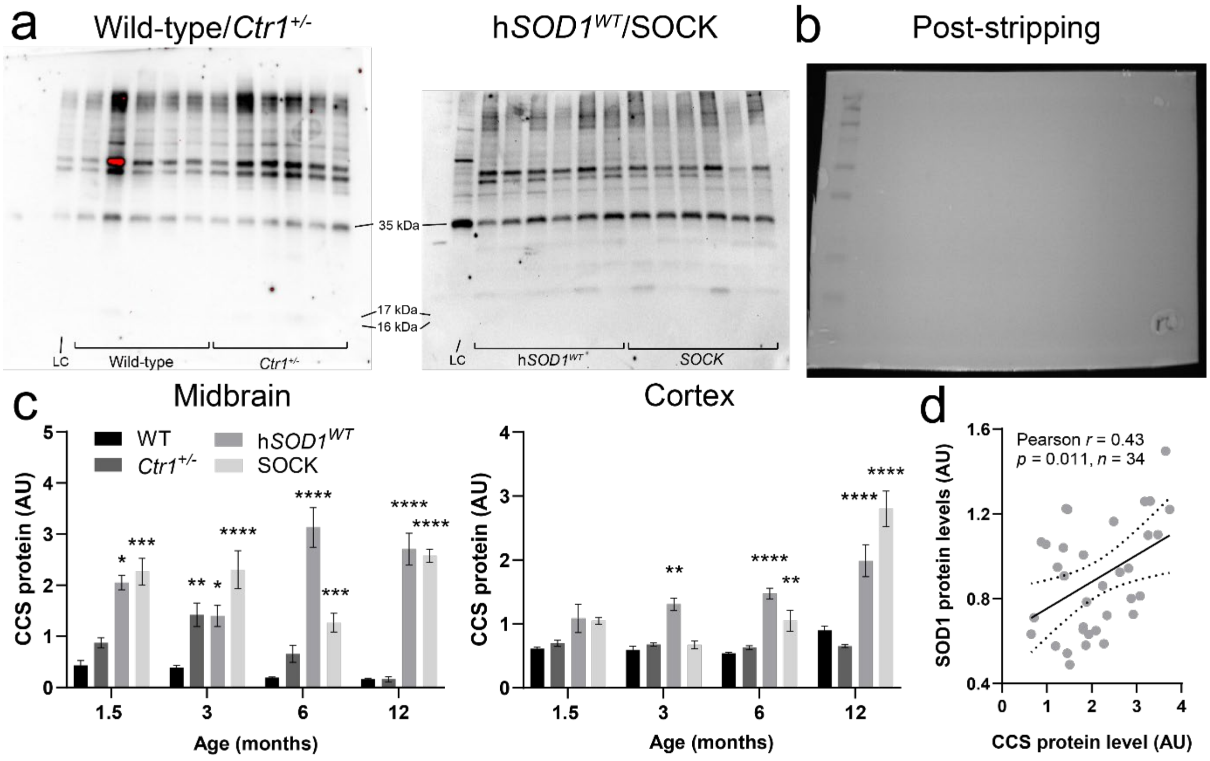


**Supplementary Fig. 5 Representative CCS immunoblots and quantification. a.** Representative immunoblots from midbrain tissue extracts, whereby samples from h*SOD1*$^{WT}$ and SOCK mice were run separately to those from wild-type and *Ctr1*$^{+/-}$ mice, given their inclusion in the same gel was impossible due to large differences in SOD1 protein expression between these groups (**Supplementary Fig. 2**). A common loading control (LC) made from pooled midbrain, cortex and liver extracts was included on all gels to enable standardization and estimation of protein expression differences between these groups. Full antibody details are listed in **Supplementary Table 8**. **b.** CCS antibody was successfully stripped prior to re-probing with SOD1 and GAPDH primary antibodies, as demonstrated by the lack of chemiluminescent signal in lanes other than the ladder. Image represents a blot that has been probed for CCS (rabbit antibody) and imaged with ECL substrate, before being stripped, washed, re-probed with HRP-conjugated anti-rabbit secondary antibody and re-imaged with ECL substrate. **c.** Quantification of CCS band density was performed in midbrain and cortex tissue extracts of all 4 mouse genotypes from 1.5 months to 12 months of age, revealing significant increases in CCS protein expression in both regions in h*SOD1*$^{WT}$ and SOCK mice across most ages examined (Two-way ANOVA with Dunnet's multiple comparisons post hoc *t*-tests). **d.** SOD1 and CCS protein levels were significantly correlated in the SOCK mouse midbrain across all ages examined, statistical test details are presented in the panel. Sample size per genotype per age: 1.5 month, $n$ = 2-8; 3 month, $n$ = 4-12; 6 month, $n$ = 10-13; 12 month, $n$ = 5-12. Data represents mean ± SEM. *$p < 0.05$, **$p < 0.01$, ***$p < 0.001$, ****$p < 0.0001$. Abbreviations: WT, wild-type.

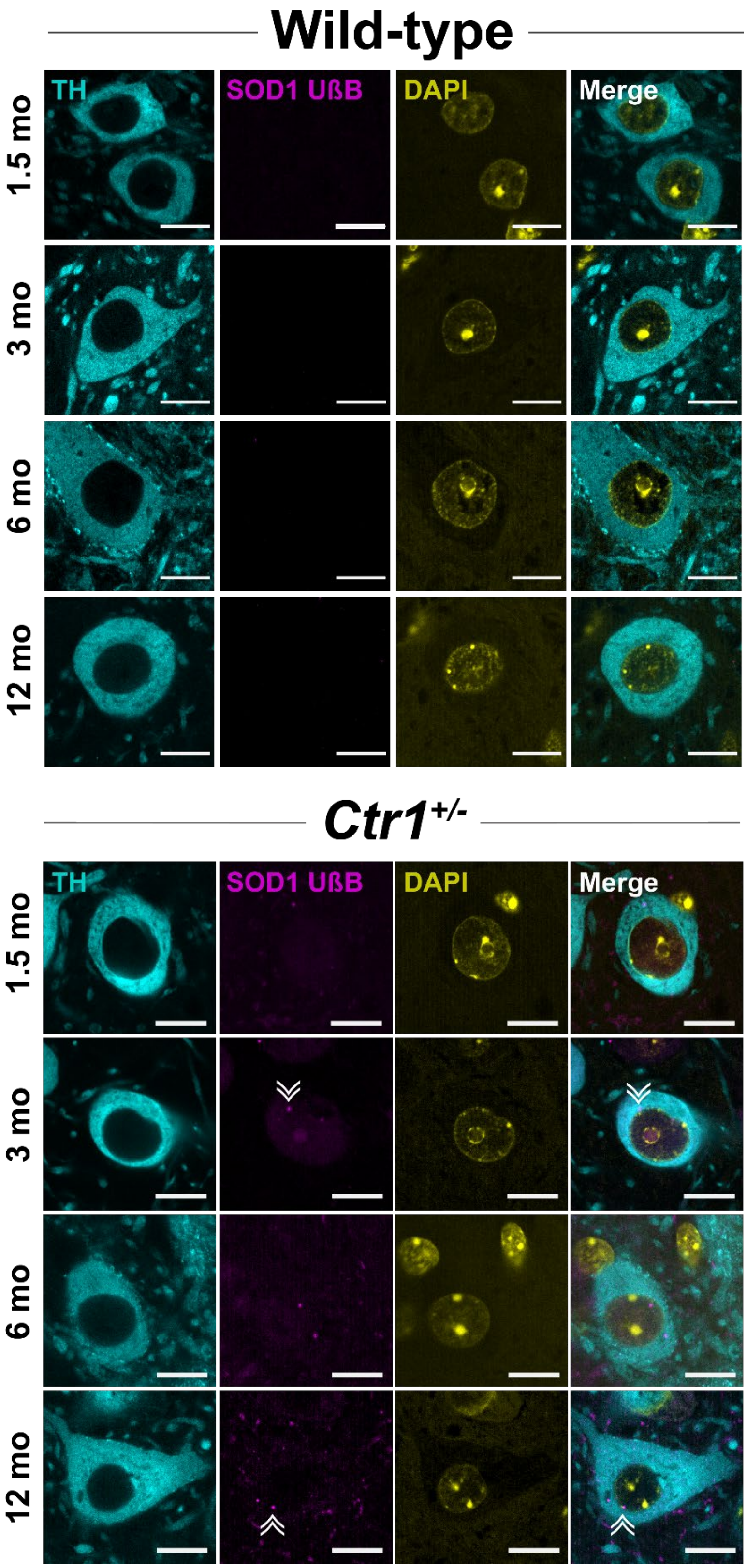


**Supplementary Fig. 6 Midbrain disSOD1 immunostaining in 12-month WT and *Ctr1*$^{+/-}$ mice**. Very little disSOD1 pathology (double white arrowheads) was detected within and surrounding tyrosine hydroxylase (TH)-immunopositive neurons (cyan) in the SNc of 12-month-old wild-type and *Ctr1*$^{+/-}$ mice using antibodies recognizing SOD1 protein in an unfolded ß-barrel (UβB) conformation (magenta). Full antibody details are listed in **Supplementary Table 8**. DAPI was used to visualize cell nuclei (yellow). Scale bars represent 10 µm.

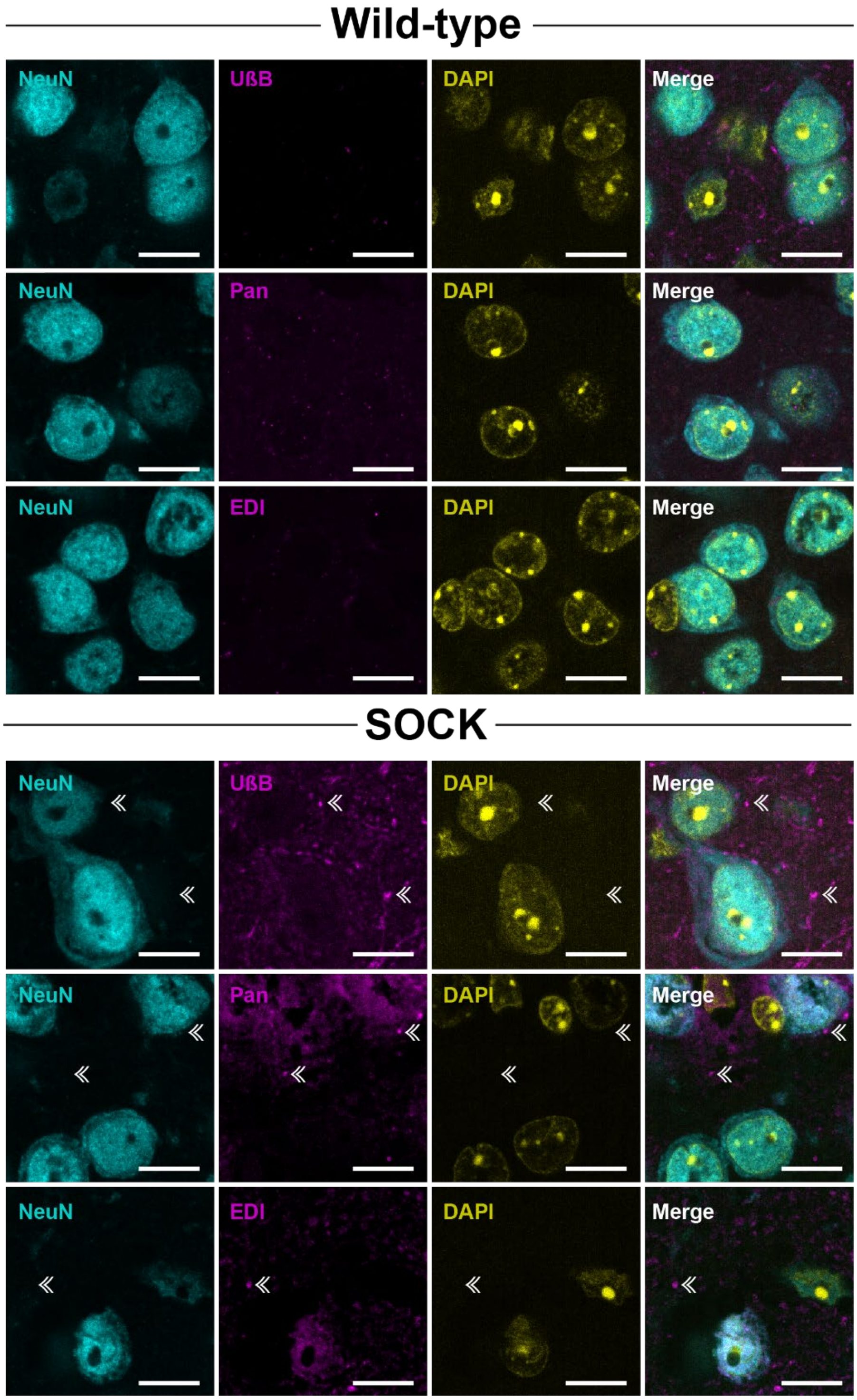


**Supplementary Fig. 7 Cortical disSOD1 pathology in 12-month SOCK mice**. DisSOD1 pathology (double white arrowheads) was detected in NeuN-immunopositive cortical neurons using antibodies recognizing unfolded ß-barrel (UβB) and exposed dimer interface (EDI) conformations of SOD1, as well as a pan-SOD1 antibody. Full antibody details are listed in **Supplementary Table 8**. DAPI was used to visualize cell nuclei. Scale bars represent 10 µm.

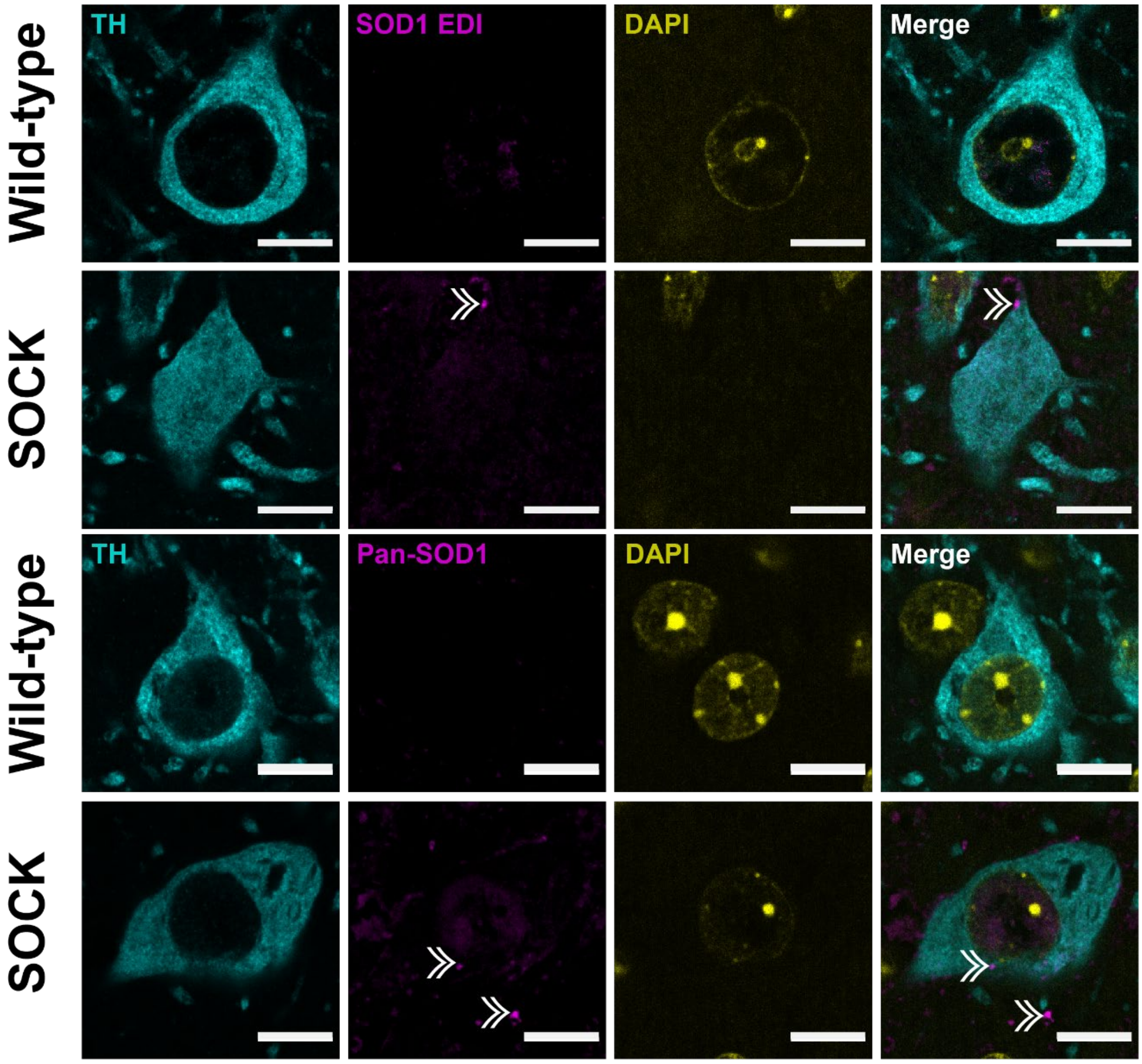


**Supplementary Fig. 8 Cross-validation of disSOD1 pathology detection in the SOCK mouse SNc**. In addition to UβB conformation-specific antibody (**Fig. 3**), disSOD1 pathology (double white arrowheads) was also detected in and around SNc dopamine neurons (tyrosine hydroxylase (TH)-positive) in the SOCK mouse brain using exposed dimer interface (EDI) and pan-SOD1 (Enzo) antibodies. Full antibody details are listed in **Supplementary Table 8**. DAPI was used to visualize cell nuclei. Scale bars represent 15 µm.

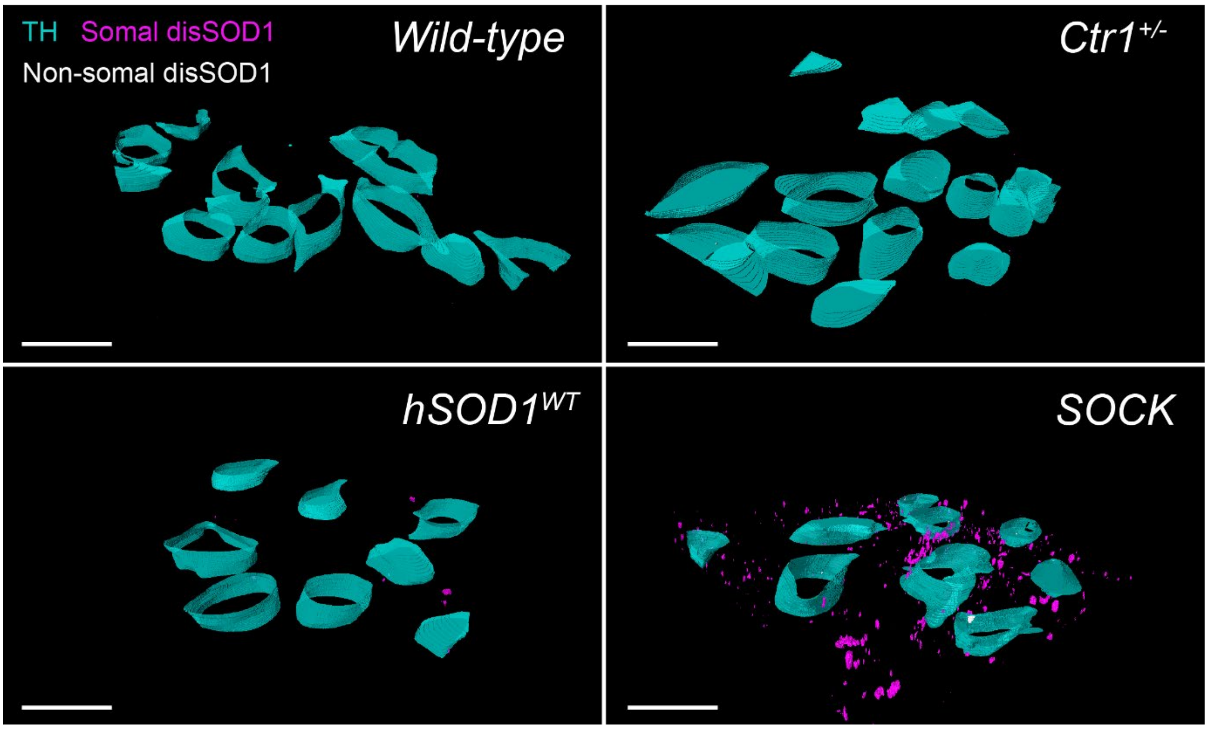


**Supplementary Fig. 9 Three dimensional reconstructions of disSOD1 aggregates in the SNc of all mouse strains.** Tyrosine hydroxylase-positive neuronal soma are highlighted in cyan, while aggregates residing within and outside of these soma are highlighted in white and magenta, respectively. Scale bars represent 30µm.

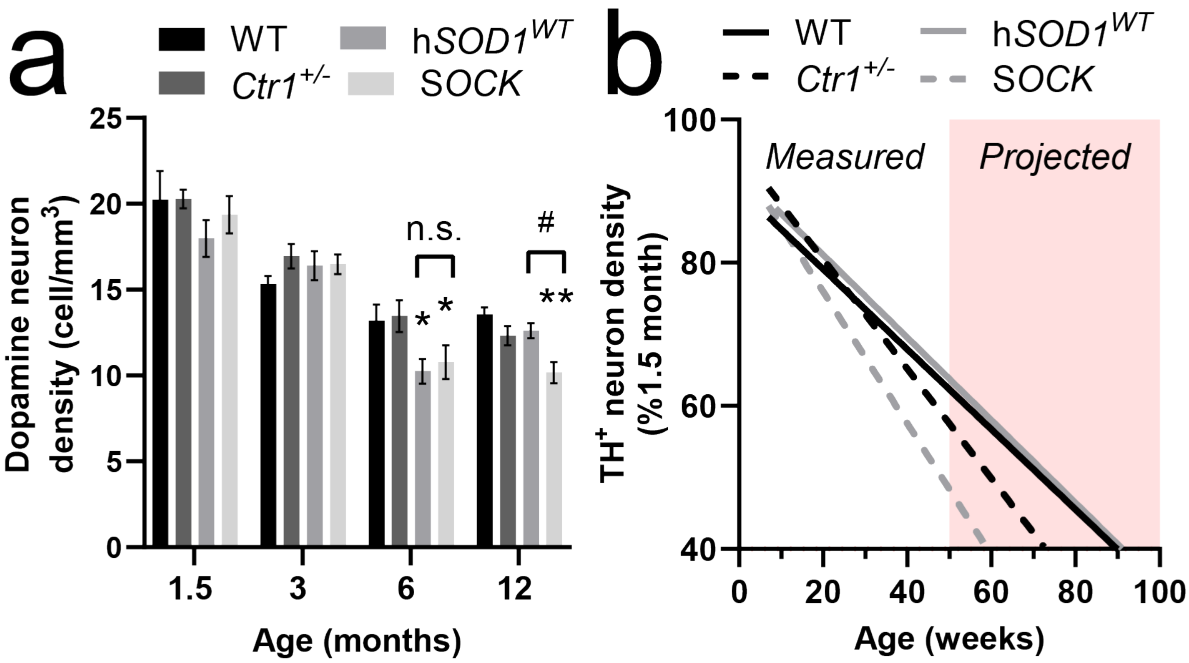


**Supplementary Fig. 10 Stereological estimates of SNc dopamine neuron density in all four mouse strains**. **a. a.** Quantitative stereology revealed significant variation in the density of dopamine neurons in the SNc of 1.5 – 12-month-old wild-type (WT), Ctr1$^{+/}$, hSOD1$^{WT}$ and SOCK mice (two-way ANOVA: age – $F_{(3, 132)} = 59.37$, $p < 0.0001$; genotype – $F_{(3, 132)} = 3.731$, $p = 0.013$), which was decreased in 6-month-old hSOD1$^{WT}$ and SOCK mice, and 12-month-old SOCK mice, compared with WT mice (Tukey's multiple comparisons post hoc test: p = 0.030, 0.041 and 0.0015, respectively). The density of nigral dopamine neurons was also significantly lower in SOCK mice compared with hSOD1$^{WT}$ mice at 12 months-of-age (p = 0.045) but not 6 months-of-age (p = 0.94). **b.** Quantified and projected stereological estimate of SNc dopamine neuron density in all mouse strains. Data represent linear regression of individual datapoints for each strain, whereby the density of TH neurons was expressed as a proportion of the average number of TH neurons quantified at 1.5 months-of-age in that same genotype. Data trends beyond 50 weeks-of-age (red zone) were extrapolated by applying linear regression to measured data, with the 95% confidence interval indicated by dotted lines.

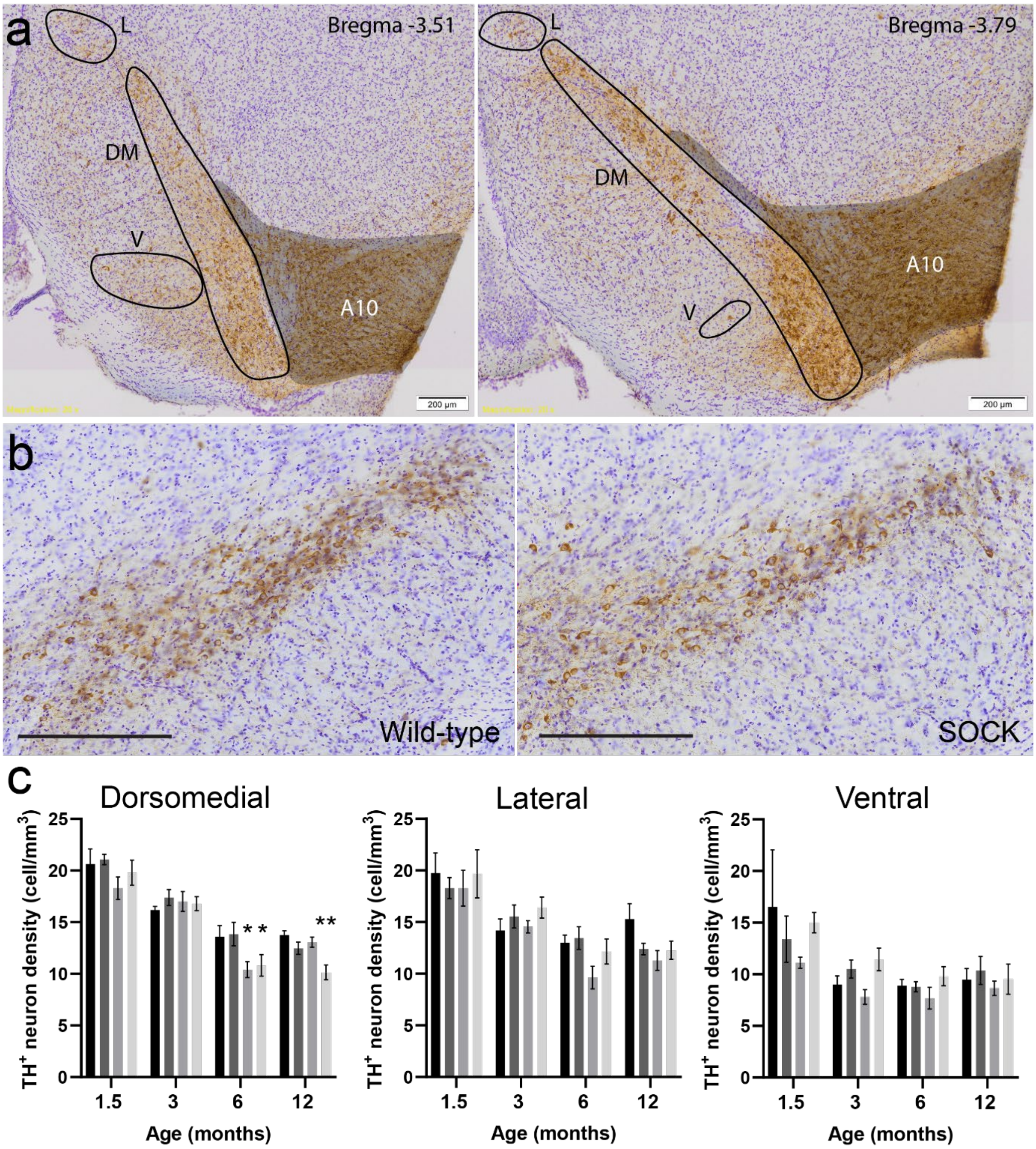


**Supplementary Fig. 11 Stereological estimates of dopamine neuron density in dorsomedial, lateral and ventral subregions of the SOCK mouse SNc**. **a.** Anatomical delineation of the dorsomedial (DM), lateral (L) and ventral (V) SNc in 50 µm midbrain tissue sections from a 12-month-old wild-type mouse. Immunostaining for tyrosine hydroxylase (TH) was performed using 3,3′-diaminobenzidine (brown) and counterstained with cresyl violet (purple). Scale bars represent 200 µm. Antibody details are listed in **Supplementary Table 8**. **b.** Representative immunostaining of the wild-type and SOCK mouse midbrain. Scale bars represent 300 µm. **c.** Differences in TH-immunopositive cell bodies in the SNc between mouse strains and ages were identified by a two-way ANOVA with Dunnet's multiple comparisons post hoc *t* tests. Data represent mean ± SEM. *n* = 3-8 (1.5 months), 5-13 (3 months), 7-14 (6 months), 8-14 (12 months) per genotype. $*p < 0.05$, $**p < 0.01$.

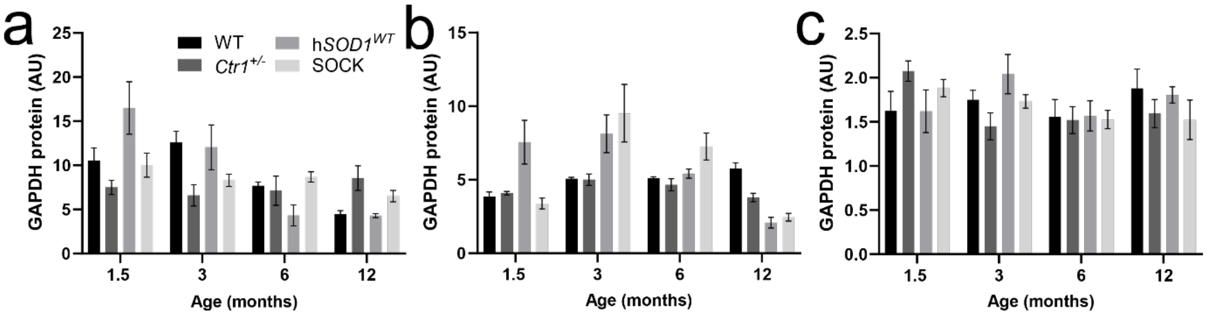


**Supplementary Fig. 12. GAPDH protein levels in the midbrain (a), cortex (b) and liver (c) of all mouse strains.** Measurements were obtained using immunoblotting, as described in **Supplementary Fig. 2,** with no variation between strains identified using a two-way ANOVA. Sample size per genotype per age: 1.5 month, $n$ = 3-8; 3 month, $n$ = 7-15; 6 month, $n$ = 10-16; 12 month, $n$ = 9-14. Data represents mean ± SEM. Abbreviations: WT, wild-type.

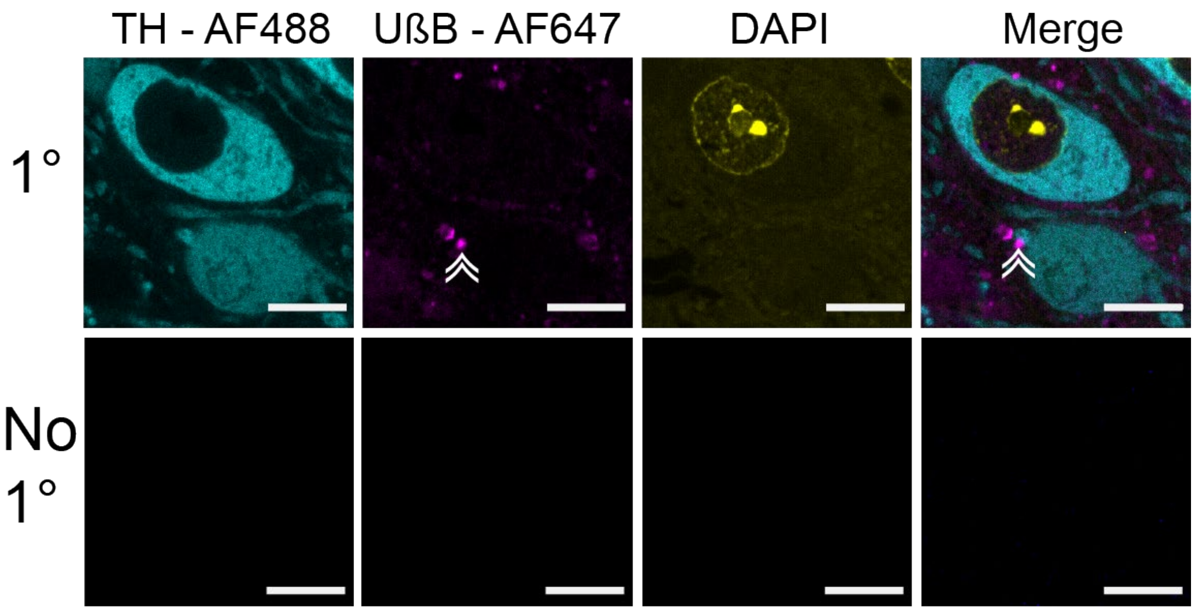


**Supplementary Fig. 13. Negative control immunofluorescent staining.** Alongside routine immunofluorescent staining (1°) for tyrosine hydroxylase (TH), unfolded beta barrel SOD1 (UβB) and cell nuclei (DAPI), we performed negative control staining (no 1°) by replacing primary antibodies and DAPI with blocking buffer. No fluorescent signals were detected when imaging no 1° staining using the same acquisition settings as those used for imaging routine 1° staining. Full antibody details are listed in **Supplementary Table 8**. Scale bars represent 15 µm.
